# Motor-driven microtubule diffusion in a photobleached dynamical coordinate system

**DOI:** 10.1101/2024.08.20.608861

**Authors:** Soichi Hirokawa, Heun Jin Lee, Rachel A Banks, Ana I Duarte, Bibi Najma, Matt Thomson, Rob Phillips

## Abstract

Motor-driven cytoskeletal remodeling in cellular systems can often be accompanied by a diffusive-like effect at local scales, but distinguishing the contributions of the ordering process, such as active contraction of a network, from this active diffusion is difficult to achieve. Using light-dimerizable kinesin motors to spatially control the formation and contraction of a microtubule network, we deliberately photobleach a grid pattern onto the filament network serving as a transient and dynamic coordinate system to observe the deformation and translation of the remaining fluorescent squares of microtubules. We find that the network contracts at a rate set by motor speed but is accompanied by a diffusive-like spread throughout the bulk of the contracting network with effective diffusion constant two orders of magnitude lower than that for a freely-diffusing microtubule. We further find that on micron scales, the diffusive timescale is only a factor of ≈ 3 slower than that of advection regardless of conditions, showing that the global contraction and long-time relaxation from this diffusive behavior are both motor-driven but exhibit local competition within the network bulk.

Significance Statement
Cytoskeletal reorganization can come at the cost of local diffusive-like disordering effects from the same active elements, but distinguishing these processes can be challenging. By photobleaching an actively contracting microtubule network, we show that the bulk redistribution of filaments exhibit a diffusion-like reorganization that can be tuned by the effective motor speed. By tuning these parameters, we show a conserved relationship between active contraction rates and effective diffusion constants, suggesting that while advection of the cytoskeletal network dominates over scales of tens to hundreds of microns, motors additionally induce a diffusive-like effect that begins to compete with advection at micron scales.

---

Whether for schools of fish evading a sea lion or in the ordered array of microtubules comprising the spindle of dividing cells, coordinated movement and emergent patterning is a hallmark of biological dynamics across all biological scales. Curiosity surrounding the underlying principles dictating such a ubiquitous feature in biology have led to an explosion of theoretical (1–4) and experimental efforts (5–9) to understand them. *In vitro* active matter systems offer a powerful means to study how cytoskeletal elements self-organize to generate a diverse array of networked structures. By mixing multimerized motors with filaments, a broad range of ordered patterns have been demonstrated, occurring in solutions which are spatially-homogeneous (5, 9, 10) or are locally defined through patterned light (11–14). A common observation from these assays is that the constitutive filaments rearrange in time under dynamics that appear to be primarily advective in nature. Recent efforts have led to several quantitative models that macroscopically describe the flow-like redistribution of microtubules under a range of conditions related to properties of the motors and filaments (15–19). In addition to advective behavior, previous theoretical studies of contractile active gels have also shown that local fluctuations within a globally contracting network can give rise to a motor-driven diffusive-like effect among filaments (20, 21), a phenomenon that has been observed experimentally (22). This seeming competition between active diffusion and advection is poorly understood and invites a rigorous approach to distinguish these two effects.

In the work presented here, we incorporate fluorescence recovery after photobleaching (FRAP) into a light-controllable kinesin motor dimerization system (12, 23, 24) to characterize the interplay of motor-driven advective and diffusive dynamics. FRAP studies have typically been accompanied by various theory-based extensions of the diffusion equation to account for convective flow, reaction of molecules, or transport (16, 25–28) and have been effectively applied to active *in vitro* systems to determine how filaments are redistributing into or elastically contracting relative to the photobleached region (21, 29–31). For our study, we photobleach a grid pattern onto a contracting microtubule network, which creates square fluorescent regions (Fig. 1(A)). By tracking the area and centroids of these regions, we are able to account for the advective contraction of the network. By measuring how the darkened photobleached lines blur, we can account for how much the microtubules in the network undergo diffusive behavior (Fig. 1(B)).

**Fig. 1.**
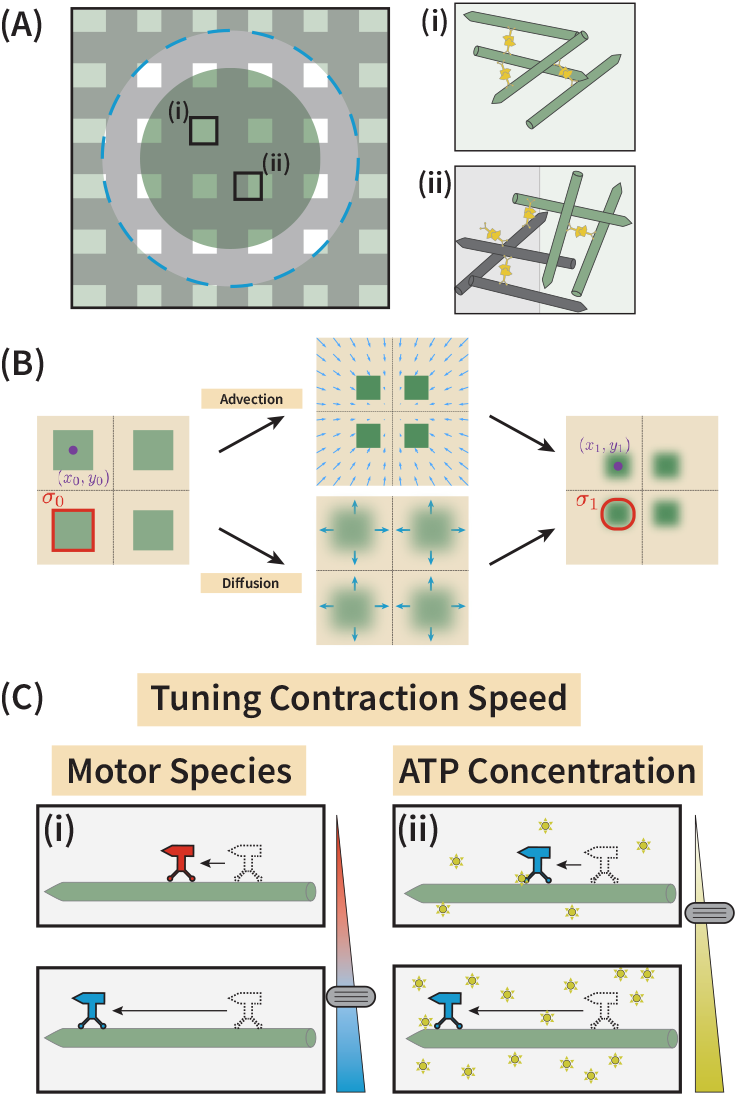
FRAP-based approaches to studying advective and diffusive-like redistribution of cytoskeletal elements. (A) Photobleaching a grid-like pattern leaves (i) squares of fluorescent microtubules (green) surrounded by (ii) non-fluorescent filaments (black) and allows us to examine the role of diffusive-like microtubule spread in the bulk of a global radially contracting network. Dashed blue circle outlines the edge of the dimerizing light inside of which the filaments couple and create a net contraction. (B) Tracking of centroids [(*x*_0_, *y*_0_) to (*x*_1_, *y*_1_)] and areas (*σ*_0_ to *σ*_1_) of the fluorescent squares allows us to quantify the advective and diffusive contributions in the contracting system. (C) The rates of these dynamics can be tuned by changing the effective motor speed through either (i) changes in the motor species or (ii) changes to the ATP concentration in the system. We tune these parameters to examine rates of contraction and bulk reorganization of microtubules in the contracting cytoskeletal network.

We find the choice of motor species (32) or the availability of ATP (33) are key parameters controlling the network dynamics (Fig. 1(C)). For example, by reducing motor speed, whether through decreased ATP concentration or slower motor species, the network globally contracts at a slower rate while the bulk of the network exhibits a decrease in effective diffusion constant. We further show that contraction rate and effective diffusion constant are linearly proportional measures across all of our conditions and give rise to a tightly bounded Peclét number slightly greater than one over micron length scales. While motors were understood to set the global contraction of the network, they play a second competing role within the network boundary that gives rise to a long-time relaxation on the local cytoskeletal structure.

## Results

### Photobleaching a grid pattern

To study the local redistribution of microtubules as the network contracts, we designed an augmented optical system that induces dimerization of kinesin through the iLid-micro system (34) and images the microtubules (12, 24). This modification includes a photobleaching element that allows us to photobleach a grid-like pattern into the microtubule channel (see Materials & Methods section and SI Sec S1.6). The end result is an array of squares roughly 12 µm in side length and 25 µm in center-to-center distance, much longer than the median microtubule length of ≈ 1.5 µm (see SI Sec S2). Fig. 2(A) shows an example of a grid pattern photobleached onto a microtubule network at different time points in its life history and the subsequent deformations of the bleached lines and fluorescent squares. As the image for the *t* = 0 sec timepoint in Fig. 2(A) shows, upon photobleaching the grid pattern, individual fluorescent squares, which we will call unit cells, are produced. Over a minute after photobleaching, unit cells contract toward the center of the network while the photobleached lines appear to blur away. By two minutes after photobleaching, neighboring unit cells appear to blend into each other and at later times any remnants of the photobleached pattern disappear.

**Fig. 2.**
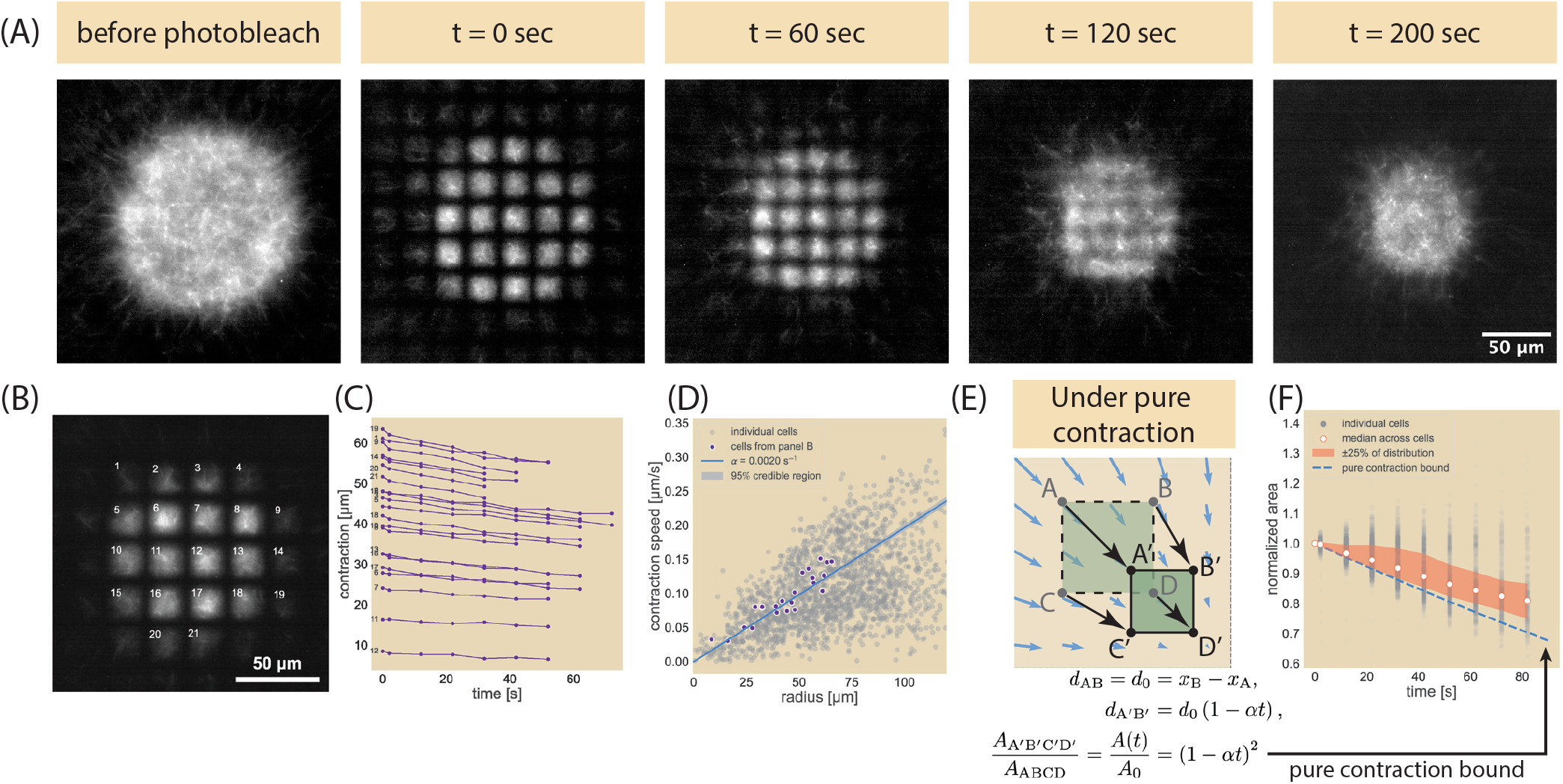
Photobleaching a grid pattern onto the contracting microtubule network. (A) Example dataset, where the microtubule field is photobleached and the deformations of the fluorescent regions observed using Ncd236 and 1.4 mM ATP. (B) Enumeration of individual fluorescent unit cells to (C) compute the distance of their centroids from the center of the network over time. Numbers correspond to labels from panel (B). (D) Plot of unit cell contraction speed as a function of their average distances from the center of the network, obtained by fitting the distance vs time data found in (C) to individual lines. The median contraction rate is *α* = 2.0 *×* 10^−3^ s^−1^ . (E) Schematic of the unit cell deformation and expected area change under a purely contracting condition. (F) The area of each unit cell is normalized against their initial area as obtained by the unit cell segmentation scheme and plotted as a function of time. The median normalized area is plotted in white among individual unit cells (gray). The red shaded region encompass points between the first and third quartiles of the distribution of all cells. Dashed blue line corresponds with the normalized area computed in (E) and using the median contraction rate obtained in (D).

### Tracking fluorescent squares shows global contraction and local diffusive spread

To better quantify and understand the global network contraction dynamics, we segmented individual unit cells and measured their centroids and areas over successive frames (see SI Sec S3 and S4 for analysis). By tracking individual unit cells such as those shown in Fig. 2(B) and computing their distance from the center of the network over successive frames (Fig. 2(C)), we can determine the local contraction speeds, where we see a rough linear correspondence between distance and time. We computed the slopes of each unit cell trajectory and compared the resultant speeds as a function of their distance from the network center (Fig. 2(D)) to find that the median contraction speed linearly increases with distance from the center, indicating a general linear contraction of the entire microtubule network. We thus fit the velocity against the radius **r** with a line passing through the origin (see SI Sec S5.1), giving the expression

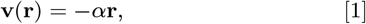

where *α* is the contraction rate and thus measure a contraction rate and 95% credible region of *α* = 2.0 × 10^−3^ ± 5 × 10^−5^ s^−1^. Data separated by experimental replicates are available in SI Sec S6.

Despite the linear global contraction observed for the centroids, a more careful examination of the unit cells reveals that the network does not simply undergo purely elastic contraction. Suppose we took two points with different x-positions but the same y-position (*x*_A_, *y*_A_) and (*x*_B_, *y*_A_), respectively, such as points A and B in Fig. 2(E) that they have a distance *d*_0_ of

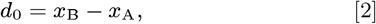

where we will take *x*_B_ *> x*_A_. If the two points were strictly subject to move from the velocity field given by Eq. 1, after time *t* their positions will have changed such that their distance *d*_1_ is now

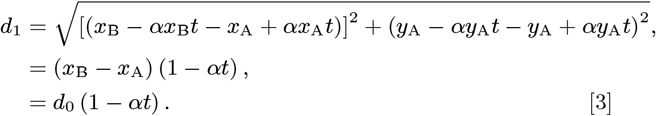

So the two points move closer by a factor of 1 − *αt* in that time. We can make a similar argument for two points vertically separated. If we imagine this for all four points that make up the corners of a unit cell (points ABDC transforming to A’ B’ D’ C’ in Fig. 2(E)) and look at the change in area, we would expect that under a purely contractile active system subject to the linear contraction rate measured from tracking the unit cell centroids, the area *A*(*t*) would change from its initial size *A*_0_ by

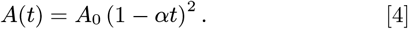

See SI Sec S7 for a more complete derivation. Fig. 2(F) shows the normalized area of each unit cell as a function of time in gray against this pure contraction scaling given as a blue dashed line. The median normalized area is shown as a white circle with a red outline. As can be seen by comparison with the shaded red region (representing the 50 percent of all cells that fall between the first and third quartiles of the distribution of cell areas), the majority of the experimental observations are above the pure contraction bound. With the area being greater than that for a purely contracting network, we conclude that despite the global contraction of the network, filaments can locally spread and reorganize in the bulk. This observation is further affirmed by the merging of originally distinct fluorescent squares in the 120-second time point of Fig. 2(A), where a purely contractile network would cause fluorescent squares to remain distinct.

### The effective diffusion constant is roughly two orders of magnitude lower than free diffusion of a microtubule

Since a purely contractile description is insufficient to fully capture the observed dynamics, we generalize our treatment of this contractile effect and accounting for diffusion using an advection-diffusion equation to model the time evolution of the tubulin concentration *c*(**r**, *t*). Such a model has a material flux **J** of the form

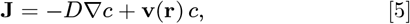

where *D* is the diffusion constant and **v**(**r**) is the velocity profile of the advective flow as a function of distance from the center of contraction *r*. As motivated by results shown in Fig. 2, we use the velocity field given by Eq. 1 with *α* as the computed contraction rate as shown in Fig. 2(D). When inserted into Eq. 5 and combined with the continuity equation, the advection-diffusion equation takes the form

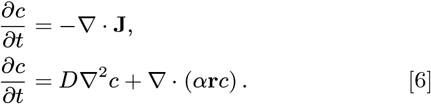

We perform a series of deep explorations of the model in SI Sec S8-S10 to better understand the time-evolution of the concentration profile subject to Eq. 6 and to validate the implementation of our finite element method (FEM) using COMSOL Multiphysics®. Ultimately, these exercises confirm the importance of using a grid-like photobleaching pattern rather than a circular pattern, the latter of which would have convoluted the contributions of the radially-directed contraction with the more-isotropic diffusive-like spread. After validating our initial simulation, we turn fully to the use of FEM simulations on Eq. 6 to model our experimental results. We perform a FEM simulation of an individual unit cell, where we fix the FEM contraction rate to be the experimentally-measured mean rate and sweep across various diffusion constants (see SI Sec S11 for implementation of a single unit cell in COMSOL). For example, for the Ncd236 motor at saturated levels of ATP, we show in Fig. 3 the family of normalized area trajectories for a single unit cell subject to the experimentally observed mean contraction rate of *α* = 2×10^−3^ s^−1^, and for diffusion constants *D* ranging from 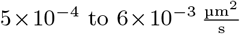 . For comparison to the experimental data, we overlay the median (circle), the 1st quartile (triangle), and 3rd quartile (plus symbol) from the distribution of measured unit cell areas trajectories, where the quartiles give a sense of the trajectory variation. Here, we see that by minimizing least squares between the experiments and the simulation conditions, the median normalized area trajectory agrees best with the FEM trajectory with an effective diffusion constant of 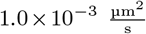 . The first quartile of area trajectories from measurements lies between the pure contraction limit where *D*_eff_ = 0 and an effective diffusion constant 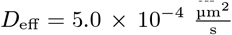. We interpret the first quartile results to mean the effective diffusion coefficient must be above 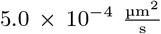 in order to model the majority of the data. The third quartile of area trajectories most closely follows the trajectory with an effective diffusion constant of 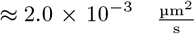 (see SI Sec S4.2 on fitting procedure), serving as a kind of upper bound. For context, the diffusion coefficients associated with the median and quartiles are all about two orders of magnitude smaller than the diffusion coefficient of a freely diffusing microtubule, which is 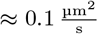 (see SI Sec S2 for microtubule length; equation for the longitudinal diffusion constant obtained from Ref (12)).

**Fig. 3.**
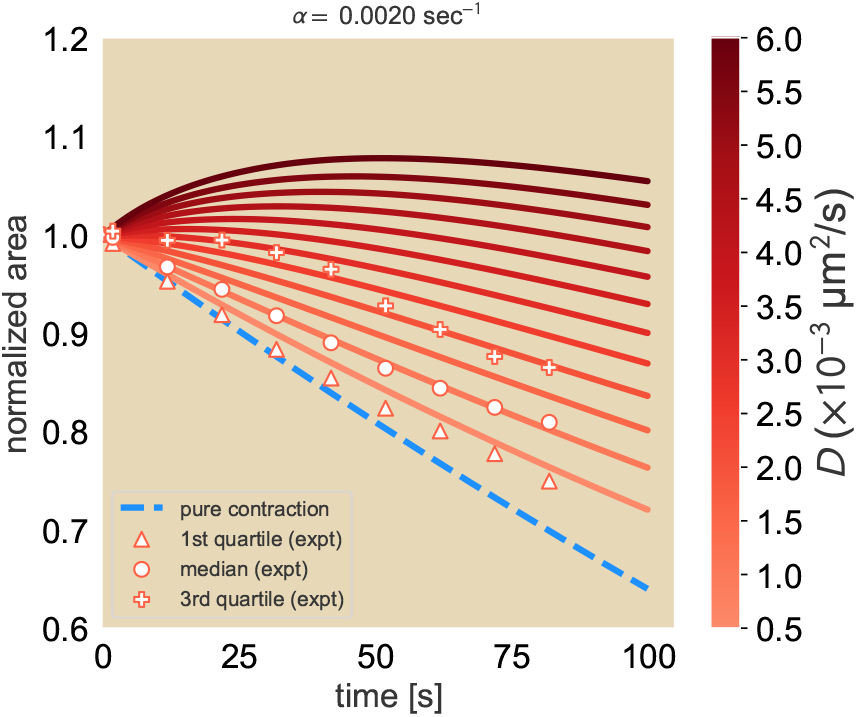
Simulated concentration profiles for a linear advection-diffusion equation. A family of curves for the expected normalized area of fluorescent squares subject to a fixed advection rate *α* = 2*×*10^−3^ s^−1^ and varying diffusion constants. The 25th percentile (triangle), median (circle), and 75th percentile (plus sign) of the area trajectories are overlayed onto the FEM results for comparison.

We also explored how well our FEM simluations could capture the qualitative features of the data set shown in Fig. 2(A), such as the merging of unit cells and the time scale of this process (see SI Sec S12). Our main finding from these efforts is, even for just qualitative comparisons, diffusion must be included in the theoretical description of the dynamics.

### Changing effective motor speed proportionally changes contraction rate and effective diffusion constant

To test whether motors play a role in the diffusive-like effect, we next tuned the effective speed of the active elements, namely, the motors themselves. Two ways in which this can be done is by the choice of motor species or by changing the concentration of ATP. We chose optogenetic versions of previously characterized motor variants that span roughly an order of magnitude in speeds: Ncd236, Ncd281 (35), K401 expressed in bacteria (12), and K401 expressed in Sf9 cells (24) (See SI Sec S14 for Ncd281 construct designs and motor speeds and processivities). Fig. 4(A) shows the motors speeds for each of these motor types and the associated contraction rate. Interestingly, we observe a roughly linear relationship between contraction rate and motor speeds. The effective diffusion coefficient also demonstrate a roughly linear trend (Fig. 4(B)). We note that even for the slowest motor Ncd281, a non-zero coefficient *D*_eff_ of 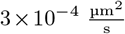 is needed to accurately describe the data. This general tendency to increase the effective diffusion constant suggests that the motor speed may be responsible for the local microtubule effective diffusion. We note both trends can be explained by just the speeds of the motor species, requiring no consideration of motor processivity.

**Fig. 4.**
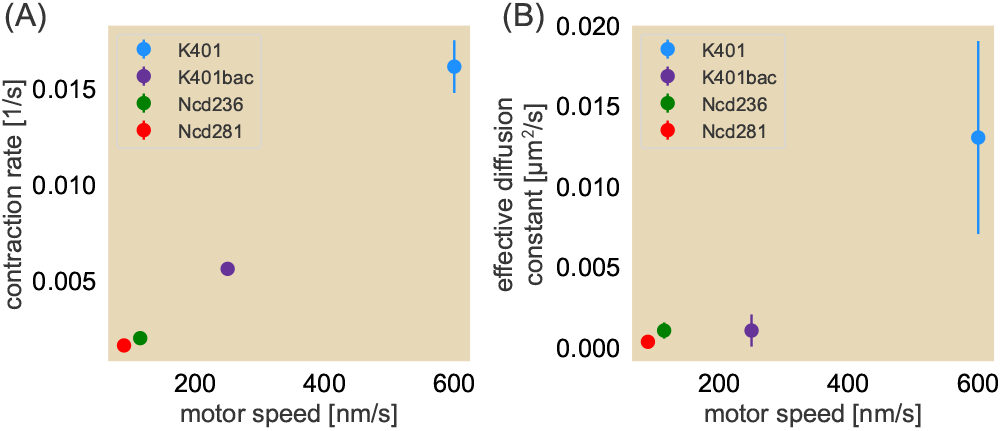
Contraction rates and effective diffusion constants for four different motor types. (A) Contraction rate as a function of motor speed. Here, the motors are (in order of motor speed as found in Table S2 of the SI Text) Ncd281 (red) (35), Ncd236 (green), K401 expressed in bacteria (purple), and K401 expressed in Sf9 cells (blue) (24). (B) Corresponding effective diffusion constants as a function of motor speed where the circles denote the medians of the experimentally obtained normalized area trajectories and error bars denote the middle 50% of the distribution. Error bars for some data points are smaller than the radius for the size of the dots.

We next examine how this local diffusive-like effect changes when we decrease the motor speed of a given species by reducing ATP concentrations. In order to traverse along microtubules, motors must hydrolyze ATP with each step they take. At saturated concentrations of ATP, motors can hydrolyze ATP at their maximal rate and therefore move at their maximum speed. At reduced ATP concentrations, the limited availability of ATP causes motors to hydrolyze ATP at a reduced rate, leading to an effective reduction in motor speed (36–38). In the context of a contracting microtubule network, we hypothesize that this decrease in motor speed translates to a reduction in contraction rate, similar to the effect observed when using a slower motor species. We further hypothesize that for a constant motor concentration, reducing the concentration of ATP will increase the fraction of motors that do not move along the microtubules and instead behave as passive crosslinkers, causing the areas of the fluorescent unit cells to fall closer to the pure contraction bound. To test this, we perform our photobleach experiment for Ncd236 and bacteria-expressed K401 at ATP concentrations spanning two orders of magnitude. We continue to use an ATP regeneration system so that the ADP concentration is negligible and therefore does not compete with ATP for the hydrolysis site (38) (see Materials and Methods). Fig. 5(A) and (C) show that for both Ncd236 (A) and K401 (C), as the concentration of ATP is decreased, the contraction rate of the network similarly decreases. At 25 µM ATP, the contraction rate with Ncd is roughly half of that at saturated levels. This concentration is also roughly the measured dissociation constant of ATP hydrolysis by the motor (39). However, at ATP concentrations below this dissociation constant, we see that the network contraction, while still occurring for an ATP concentration of half the dissociation constant for Ncd236, dramatically slows down. We fit the contraction rate against ATP concentration to the best fit of a Michelis-Menten equation to find that the expected dissociation constant for ATP hydrolysis in the contracting network context (Ncd236: 30 ± 13 µM; bacterial-expressed K401: 47 ± 13 µM) is roughly the same as for measured motor speeds (Ncd236: ≈ 23 µM (39); bacterial-expressed K401: 28.1 ± 0.9 µM (38)).

**Fig. 5.**
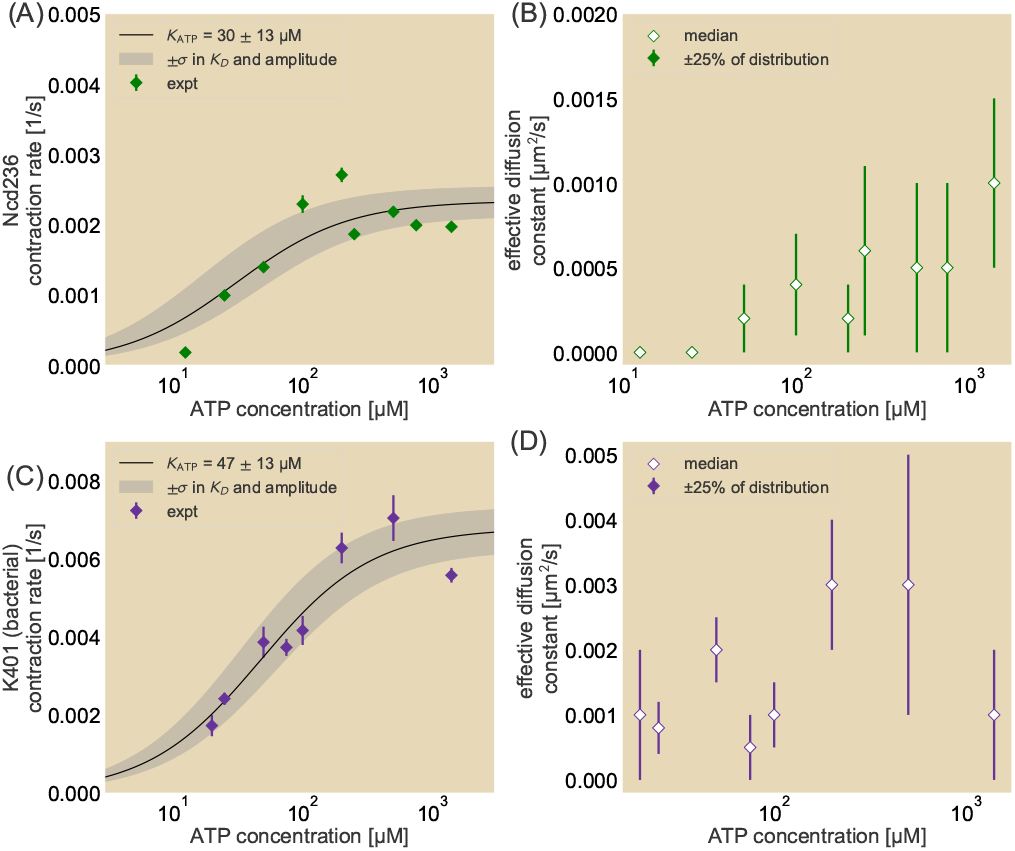
Contraction rates and effective diffusion constants over a range of ATP concentrations. Contraction rates (A and C) and effective diffusion constants (B and D) as a function of ATP concentration in the system. Motors used are Ncd236 (A and B) and K401 expressed in bacteria (C and D). Black line represents best fit to a Michaelis-Menten equation. Edges of the gray shaded region bounded to the left (right) using the Michaelis-Menten equation where the amplitude is the best fit plus (minus) one standard deviation and the dissociation constant is the best fit minus (plus) one standard deviation. Effective diffusion constants fitted to the median area trajectories with error bars corresponding to fits spanning the middle 50% of the distribution motor types are presented.

Decreasing the ATP concentration overall reduces the effective diffusion constant for Ncd236 (Fig. 5(B)). As the network dynamics scale with ATP concentration in a similar way to single-motor kinetics, our results show that motors are drivers of not only the contraction rate but also the local diffusion-like relaxation of the network. On the other hand, such a trend is less clear for bacteria-expressed K401 at some ATP concentrations (Fig. 5(D)), suggesting a need to further explore the underlying cause for this behavior.

### Contraction rate and effective diffusion constant are unified in the Péclet number

From tuning the motor type and ATP concentration in our *in vitro* kinesin-microtubule system and measuring the resultant contraction and diffusion rates, we see when parameters increase the contraction rate of the network they also similarly increase the effective diffusion constant. This suggests a relationship between the advective and diffusive properties.

To characterize this, we make use of the Peclét number, a non-dimensional ratio between the rates of advection and diffusion in the system, given by

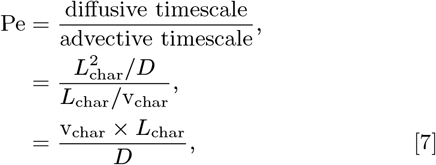

where *L*_char_ and v_char_ are the characteristic length scale and characteristic advective velocity, respectively. *L*_char_ determines the length scale in the system over which the advective and diffusive timescales are compared. In the system, candidates for *L*_char_ may be as small as a typical microtubule in the assay and as large as the size of the contracting system. As the microtubule length is within an order of magnitude of the photobleached line and gives a sense of the local competition between diffusion and advection, we choose *L*_char_ = 1.5 µm. We relate v_char_ to *L*_char_ through the global contraction rate of the network *α*. In other words, v_char_ = *α × L*_char_. So we have

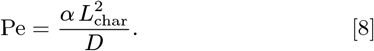

To estimate the Peclét number, we plot the median contraction rates multiplied by the square of a characteristic length scale against the corresponding effective diffusion rates for all experimental conditions (ATP concentration as diamonds and motor species as hollow circles) in Fig. 6, which demonstrates a roughly linear relationship. The slope of the line through a set of constants gives Pe, specifically Pe = 2.6 ± 0.2. To get a sense for how much Pe varies due to variability within conditions, we find Pe_25_ = 4.5 ± 0.5 and Pe_75_ = 2.4 ± 0.1 for the first and third quartile datasets, respectively (see SI Sec S14), suggesting that Pe is tightly constrained. In all cases, Pe ≳ 1, suggesting that the effect of diffusion is smaller than that of advection, but comparable to within an order of magnitude. This makes sense as the net effect of the unit cells, despite exhibiting a local diffusive-like effect, shrinks in area. Further, the rather narrow range in the Péclet number despite the spread in quartiles further suggests that the speed of the active elements sets both the global contraction of the network and the local spread of individual filaments. We recall that our choice of *L*_char_ allows us to examine the local competition between advection and diffusion. Had we chosen the side length of a unit cell or the size of the system as our size scale, we would see the increase in *L*_char_ results in Pe ≫ 1, demonstrating the greater dominance of the advective component over larger length scales, consistent with the net contraction in the network. Our results indicate that while advection generally dominates over diffusion, most notably over longer length scales, the close linear relation between the two rates suggest that they are both set by the speed of the motors.

**Fig. 6.**
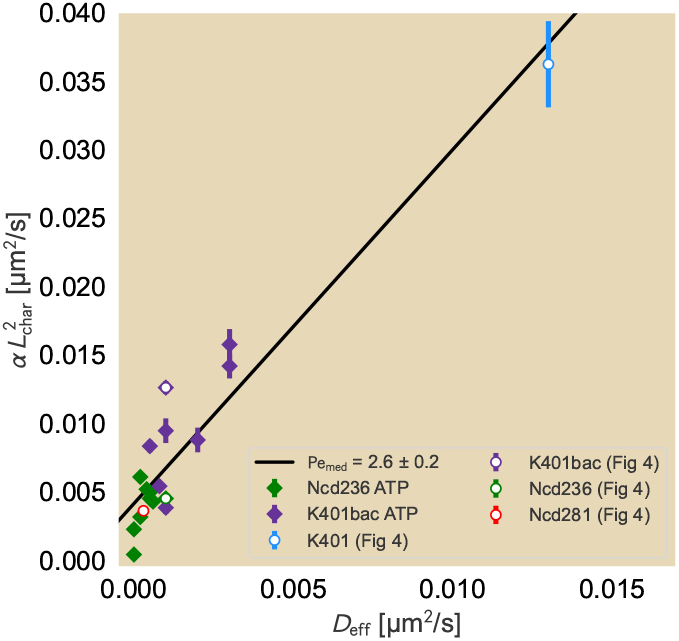
Relation of contraction rate and effective diffusion constant. Comparisons of contraction rate to effective diffusion constant are made for effective diffusion constants fitted to the median normalized area trajectories and obtained across all ATP concentration (diamonds) and motor species (hollow circles with colors matching those in Fig. 4) conditions. Contraction rates are multiplied by the square of a characteristic length scale, in this case roughly the median length of a microtubule in experiments (1.5 µm), to match the units of the effective diffusion constant. Slopes of lines are best fits of Pe, which are reported with their respective standard deviations in the legend.

## Discussion

The dynamic cytoskeleton is critical to carrying out key processes within cells, such as the formation and maintenance of the mitotic spindle (40), cell division by cytokinesis (41), and as centers of morphogenetic information (42). Such motor-filament structures are vital to a cell or organism, but how the constituent cytoskeletal elements reorganize to reach the same end configuration due to changes in biochemical conditions has been unclear. In order to understand this response by the kinesin-microtubule network, we developed an experimental framework for probing the bulk redistribution of the filament network using a grid photobleaching pattern. By photobleaching the network, we observe that microtubules will undergo a diffusive-like spread that locally opposes the global, linear contraction of the system. The diffusive-like behavior and contraction rate are jointly tuned by changes to the effective motor speed either from using different motor species or altering the ATP concentration. These effects also appear to occur from tuning crowding agents, a topic that we discuss in SI Sec S15-S16. In short, not only is the contraction an actively driven phenomenon, but so too is the diffusive-like behavior.

As we observed a general increase in effective diffusion constant with the increase in contraction rate, we further probed this relationship to find a roughly linear relationship between the two measurements. This strongly suggested that the Péclet number, which is the ratio of the diffusive timescale to the advective one, remains roughly constant within the bulk of the contracting network regardless of the biochemical conditions we used here and further suggests that motor velocity not only sets the rate that the system contracts but also the effective diffusivity in the bulk.

Active diffusion has been observed in other active systems both *in vitro* (21, 29–31) and *in vivo* (43–45). This phenomenon is particularly exciting in those cases where the diffusive-like motion exhibits a dependency on the availability of ATP. *In vitro*, a Michaelis-Menten like relationship has been observed between ATP concentration and the spatial rate of deformation of the system (30). This relationship between the rate at which deformations occur and ATP concentration is consistent with our observations that both the macroscopic deformation rate and local diffusion is determined by an effective velocity of the motor. Furthermore, the fact that this ATP dependence also occurs in *in vivo* systems (44, 45) suggests interesting implications in the ability of cells to carry out key enzymatic reactions. One key parameter that has not yet been explored is the competition of ADP in the system. We recall that our work utilized an ATP recycling system that allows us to ignore such competing effects. However, cells also have a supply of ADP that may compete with ATP for the ATP hydrolysis site of motors (38). A natural extension of this study would involve systematically tuning the ratio of ATP to ADP to observe the effects of reorganization due to this competition and may provide key insights on the role of metabolic activity on altering the rates of ATP-dependent processes.

Various quantitative models have been made to recapitulate the experimentally observed motor-filament dynamics (11, 18, 19, 21, 23, 46). These models incorporate contractile stresses in the system, either through motor-driven activity or Stokeslets in the flow field, resulting in a net elastic behavior (11, 19, 23). As we report here, a long-time relaxation term is required to create the diffusive-like filament redistribution in the network bulk. Recent treatments to introduce a long-time relaxation, as found in Ref (47), propose force-balance approaches that incorporate a viscoelastic stress to locally oppose the active stress underlying the elastic contraction. As our findings indicate that the relaxation is motor driven and transports filaments based on the direction of their orientation, we propose that an active viscoelastic-like stress term would be necessary and would need to depend on the motor speed and local polarity of the network. Indeed, other works account for the orientation 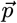 of the filaments not only for driving the movement of motors but as time-dependent variables through crosslink-generated torques (21, 46), offering encouraging pathways to recapitulate active cytoskeletal reorganization.

Our work has provided deeper insights into the extent of filament redistribution during network reorganization, where the active element not only drives the global contractile behavior but also generates a local redistribution that can be tuned by their effective speeds. Our findings leave many exciting questions about these self-organizing systems. Much is still not known about the origins of the network formation from the initially random orientation and uniform distribution of filaments prior to contraction. Specifically, the key criteria of the formed network, whether in the form of a density or order dependence, to drive the contraction process remain unclear. Photobleaching as applied in our work here provides a helpful macroscopic view of filament reorganization that can serve as a complement to other methods that are likely required to probe the dynamics of the filaments in the network, such as their orientation when they become coupled by the multimerized motors. Furthermore, the work presented here offers motivation for examining redistribution of molecules in other actively driven contexts, such as in systems of opposing motors or subject to more complex iLid-micro activation geometries (23) and with the introduction of ADP to compete against ATP-dependent reactions.

## Materials and Methods

### Microscopy set-up

The microscopy elements used to activate the iLid-micro dimerization and image the different fluorescence channels are similar to those found in Ross *et al*. (12). Briefly, a digital light processing projector from Texas Instruments was used to activate the motor dimerization and image the microtubule channels. An excitation filter wheel was placed in front of the projector to filter out the different channels. Photobleaching was performed using a diode laser with a center wavelength of 642 nm. A piezoelectric mirror gimbal mount from Thorlabs was placed downstream of the laser to deflect the beam path over a small range before the laser light passes through a cylindrical lens array inserted into a direct-drive rotation mount. The gimbal mount can then sweep the projected lines laterally to thicken the photobleaching lines before the rotation mount is rotated 90^°^ and the gimbal mount changes the deflecting angle of the beampath in the orthogonal direction. Imaging is performed using a 20x objective. More details are available in the SI Sec S1.6.

### Microtubule network assay

The microtubule network formation and contraction assay is set up similarly as in Ross *et al*. (12). Micro- and iLid-tagged motors are mixed in equal motor monomer ratios with GMPCPP-stabilized microtubules labeled with Alexa 647 in a reaction mix containing among other components ATP, ATP recycling reagents including pyruvate kinase and phosphoenolpyruvate (PEP), and pluronic as a crowding agent. While elements of the oxygen scavenging are kept in the reaction, the glucose oxidase is removed from the reaction to ensure photobleaching. Removal of these oxygen scavengers minimally affects fluorescence intensity during imaging from using the projector over the time range over which the data is analyzed, as shown in SI Sec S3.

### Image acquisition arrangement

Control of the light-dimerizing activation, photobleach laser activation, and imaging are performed through the Micro-Manager (MM) software (48, 49) while photobleaching is synchronized using a series of in-house compiled executable files that control the movement of the gimbal and rotation mounts. During acquisition, a beanshell script in MM changes the projection pattern on the DLP to create a circular light pattern for the iLid activation and full field for the imaging channels. When the desired state of the microtubule network is reached for performing photobleaching, the script completes the image acquisition cycle before turning on the photobleaching laser and calling to the executables to create the grid before the next acquisition cycle.

### Motor purification

Kinesin motors are expressed using the pBiex-1 vector transfected in Sf9 suspension cells. Cells are transfected at 5-7 µg for every 15×10^6^ cells at a starting concentration 10^6^ cells per mL of Sf900-III media using a liposome-based transfection regent (Escort IV Transfection Reagent). Cells are harvested ∽60-72 hours after transfection and purified using the FLAG affinity tag and anti-FLAG antibody resins. Proteins are stored in 50% glycerol by volume with 1.5 mM DTT, 50 µM EDTA, 50 µM EGTA, and 15 µM ATP and stored at -20^°^C. Full storage buffers and final concentrations of components are available in the SI.

## Data Availability

All data and code are publicly available. Raw image files and COMSOL simulation file can be downloaded from the CaltechDATA research data repository under the DOI:10.22002/f23ds-f2v87. Analyzed data files and code generated by Python (for analyses) and BeanShell Scripts and C# (for hardware communication) for the work presented here are available on the dedicated GitHub repository under the DOI:10.5281/zenodo.12806576.

## ACKNOWLEDGMENTS

We thank members of the Rob Phillips lab for useful discussions. We would also like to thank the David Van Valen and Rebecca Voorhees labs for providing resources for performing the protein expression and purification. We also thank Justin Bois, Griffin Chure, Peter Foster, Sebastian Fürthauer, Victor Gomez, Stephan Grill, Catherine Ji, Frank Jülicher, Matthias Merkel, Daniel Needleman, Leïla Perié, Henk Postma, Madan Rao, Shahriar Shadkhoo, and Fan Yang. This work was supported by 1R35 GM118043 and 2R35 GM118043 Maximizing Investigators’ Research Awards (MIRA) (to R.P.). S.H. was also supported by the Foundational Questions Institute (FQXI) (R.P.).

