## Supplementary Information for "Motor-driven microtubule diffusion in a photobleached dynamical coordinate system"

**S2 Microtubule length extraction** **5**

**S3 Image processing: global drift correction** **6**

**S4 Quantifying microtubule unit cell dynamics** **7**

**S6 Experimental variation of contraction speed and normalized area trajectories** **16**

**S7 Deformation of a square due solely to contraction** **19**

**S8 2D Linear Advection-Diffusion Model** **21**

**S12 FEM results of advection-diffusion equation on a simulated unit cell array** **33**

**S13 Motor Constructs** **34**

**S14 Variability in Peclét number** **36**

**S15 Computing depletion forces** **36**

**S16 Microtubule bundling can affect both contraction speed and filament redistribution** **41**

### S1 Materials and Methods

#### S1.1 Motor purification

Plasmids containing the gene encoding the motor-fluorescent protein-light-activated dimerization-FLAG tag construct with the pBiex-1 vector are transfected in Sf9 suspension cells for 60-72 hours at 27°C on shakers rotating at 120 rpm. Cells are then lightly centrifuged at 500 rpm for 12 minutes to remove the supernatant before resuspending in lysis buffer (100 mM NaCl, 2 mM $MgCl_2$, 0.25 mM EDTA, 0.5 mM EGTA, 0.25 % Igepal, 3.5% sucrose by weight, 10 mM imidazole pH 7.5, 10 µg/mL aprotinin, 10 µg/mL leupeptin, 1 mM ATP, 2.5 mM DTT, and 0.5 mM PMSF) and leaving on ice for 20 minutes. Cells are then spun down for 30 minutes at 50k rpm after which the lysate is transferred to tubes containing mouse monoclonal anti-FLAG resin (Sigma A2220) and slowly rotated at 4°C for 1.5∽3 hrs to allow protein binding to the resin via the FLAG tag. Resin-bound protein are washed three times by spinning down at 2000× $g$, clearing the supernatant, then resuspending by tube inversion in wash buffer containing 15 mM KCl, 0.5 mM, 0.1 mM EGTA, 0.1 mM EDTA, 2 mM imidazole pH 7.5, 10 µg/mL aprotinin, 10 µg/mL leupeptin, 0.3 mM DTT, and ATP in 3 mM, 0.3 mM, and 0.03 mM concentrations for the first, second, and third washes, respectively. After the third wash, the protein are spun down again at 2000× $g$ and most of the supernatant is removed, leaving the resin bed and roughly an equivalent amount of supernatant by volume in the tube. The resin bed is resuspended and FLAG peptide (Sigma F4799 or Thermo Scientific A36805) is added at a final concentration of 0.5 mg/mL before rotating for 3 hrs at 4°C. After incubating to allow the peptide to outcompete the protein for resin binding, the protein are spun down again at 2000× $g$ with the supernatant extracted and further spun down using centrifuge columns with ∽30 µm pore sizes to further separate proteins from any collected resin beads. Flow-through of clarified protein are spin concentrated using a 50 kDa filter tube to a final concentration of 2-2.5 mg/mL before diluting in 100% glycerol of the same volume for storage.

#### S1.2 Stabilized microtubule polymerization

Fluorescently labeled stabilized microtubules are prepared as in [1, 2]. After flash thawing at 37°C and kept on ice, a combination of ≈ 1.5 mg unlabeled and 100 µg labeled tubulin are diluted to 7.5 mg/mL and 0.5 mg/mL, respectively, in M2B 6.8 containing DTT and GMP-CPP at final concentrations of 1 mM and 6mM, respectively. The tubulin mixture is then incubated on ice for 5 minutes in an ultracentrifuge tube before ultracentrifugation at 90,000 rpm at 4°C for 8 minutes. Avoiding the pellet at the the bottom, the supernatant containing tubulin monomers are then placed in a new Eppendorf tube and incubated at 37°C for 1 hour, typically in a water bath, during which the tubulin is polymerizing and stabilizing with GMPCPP. The microtubule mixture is then aliquoted into individual PCR tubes while constantly being suspended in the mixture by stirring with a pipette tip. PCR tubes are then briefly spun down with a tabletop minicentrifuge before flash-freezing with liquid nitrogen and placed in a -80°C freezer for long-term storage. Microtubules are then prepared for experiments by immersing the PCR tube in 37°C water immediately when taken out of the freezer to quickly thaw.

#### S1.3 Glass slide treatment

Corning glass slides and No. 1.5 Deckgläser coverslips are coated with an acrylamide solution to prevent the adhesion of proteins from the light-dimerized activation assay to the surface. The acrylamide coating is done similarly to that demonstrated in [3]. Prior to application of the solution, slides and coverslips are separated by placement in appropriately sized containers and rigorously cleaned through a series of solutions and sonicating. First, slides are immersed in 1% Hellmanex to remove dirt particulates, sonicated, repeatedly rinsed with deionized water (DI $H_2O$), then repeatedly rinsed with ethanol. Slides are then sonicated in 200 proof ethanol before rinsing again with DI $H_2O$. After rinsing, slides are sonicated in 0.1 M KOH and subsequently rinsed in double-distilled water (dd$H_2O$). Finally, trace metals are removed by immersing in 5% HCl for 4 hours. After repeatedly rinsing in dd$H_2O$, slides are stored overnight with MilliQ ultrapure water.

Upon cleaning and before the acrylamide coating, a silane solution is made first by mixing 98.5% 200 proof ethanol and 1% acetic acid before adding 0.5% trimethoxysilyl methacrylate and immediately pouring into the containers holding the slides and coverslips. After roughly 30 minutes, slides are rinsed twice in 200 proof ethanol before drying with $N_2$ air and baking at 110°C for 10-15 minutes to cure silane onto surface with oxygen bonding.

The polyacrylamide solution is made by mixing 950 mL dd$H_2O$ with 50 mL 40% acrylamide and degassing under vacuum for 30 minutes. The solution is then under constant mixing on a stir plate with a stir bar during which time 350 µL TEMED and 700 mg ammonium persulfate (APS) are added to the solution. The acrylamide solution is immediately added to the slides and coverslips and incubated overnight. Slides are placed in 4°C for long-term storage.

#### S1.4 Flow cell chamber preparation

Flow cells for all light-dimerized activation assays are prepared by thoroughly rinsing an acrylamide-coated glass slide and coverslip in dd$H_2O$ and air drying with $N_2$ gas. A piece of parafilm with three channels each cut 3 mm wide is placed on the glass slide with the long axis of the channels running along the length of the slide. The coverslip is placed on top of the parafilm with pressure applied to flatten out the film. The flow cell is then briefly placed on a hot plate set at 65°C to warm the parafilm, allowing extra pressure on the contact points between the film and the glass to better seal the chambers.

#### S1.5 Light-dimerized activation assay preparation

Photobleaching experiments require an energy mix to maintain stability and function of microtubules and motors while constantly supplying kinesin motors with ATP to contract the microtubule network. This energy mix is slightly altered from that used by Ross *et al.* [1] with the major changes being a change in acidity for K-PIPES from pH 6.8 to pH 6.1 and the absence of gluocose oxidase to allow for photobleaching. iLid- and micro-tagged motors with the same fluorescent protein are each added to the reaction mixture at final concentrations of 40-100 nM with stabilized microtubules added at a final concentration of 1.5-2.5 µM tubulin. Concentrations of motors and tubulin are tuned to ensure that the microtubule network 1) contracts into an aster, which can fail to occur with too few motors or tubulin, and 2) without an influx of microtubules from outside of the light-activation region, which can occur from too much tubulin or too many motors dimerizing in the absense of light.

#### S1.6 Optical set-up

The sample is imaged and photobleached using a super planar fluorescence 20x objective from Nikon (numerical aperture 0.45). Image acquisition is performed using a FLiR Blackfly monochrome camera (BFLY-U3-23S6M-C) with three filters in front of it: a Semrock Brightline dual-band pass filter centered at 577 nm (28.3 nm FWHM bandwidth) and 690 nm (55.1 nm FWHM bandwidth); and a

Semrock StopLine single-notch filter at 532 nm (17 nm notch bandwidth) to suppress transmission of the YFP laser to the camera.
Activation of motor dimerization and imaging of the microtubules is performed using a digital light projector DLP Lightcrafter Display 4710 EVM Gen2 from Texas Instruments. The DLP projects white light while a motorized filter wheel sets the transmissible range of wavelengths onto the sample (beam blocker for no light, 460/50 nm filter for blue light for iLid-micro dimerization and 630/38 for microtubule imaging). Photobleaching of microtubules is performed using a 645 nm laser. The laser path is set to pass through a cylindrical lens array that transforms the collimated light pattern into a series of lines along one axis. The cylindrical lens array is mounted onto a rotation mount to allow for photobleaching of vertical and horizontal lines to generate the grid pattern. To ensure that the photobleached lines persist for multiple frames of the image, the laser passes through a gimbal-mounted mirror that deflects the beam over a small range of angles. By deflecting the laser light off of the mirror through two lenses with the same focal length $f$ and a second, stationary mirror placed $4 \times f$ away from the gimbal-mounted mirror before passing the laser through the cylindrical lense array, the transformed laser lines can be swept out. We use this beam steering approach to photobleach thicker lines.

To perform the activation and imaging patterns, we supply µManager with TIFF image stacks of matching pixel dimensions as the projector and use a Beanshell script modified from Ross *et al.* [1] to use the correct TIFF image in the stack. The TIFF stack contains a blank image (all pixel values 0) for when the laser is turned on (which is also used in conjunction with the beam blocker to prevent light from passing onto the sample outside of the activation and imaging cycles); a maximum pixel intensity image for the microtubule imaging, and a circular pattern in a blank background for the circular iLid-micro dimerization activation pattern. The primary modification to the Beanshell script is the incorporation of a timer for when the photobleaching will be performed. Once the experiment reaches the desired time, the imaging pauses while the Beanshell script turns on the laser and executes a series of custom written executables that sweep out the laser lines to create thicker parallel photobleached lines, turn off the laser, rotate the cylindrical lense array, then reactivate the laser and sweep out the laser lines in the orthogonal direction to generate the grid pattern. Upon finishing this command, the laser is shut off and imaging resumes. The entire photobleaching is performed within a roughly 10-15 second window.

#### S2 Microtubule length extraction

Stabilized microtubules imaged under total internal reflection fluorescence (TIRF) microscopy such as the ones shown in Fig. S1(A) were analyzed similar to that discussed in [1] in order to extract their lengths. Briefly, due to the uneven illumination that can occur in the image, images were first background corrected using a local thresholding method known as Niblack thresholding [12] with window size of 3 pixels and $k$ value of 0.001, which determines how many standard deviations below the mean pixel value that one sets the cut-off in the window. Although the array is a series of pixel values to be weighed against the original image, we found that this array already improved the image contrast. With this improved contrast but considering that the result is still a nonbinary image, we used Otsu thresholding on the Niblack theshold array to extract the microtubules from the background. The result is shown in Fig. S1(B).

Using the binary image which contains extracted microtubules, we imposed a morphological closing algorithm to reconnect any microtubules that were broken during the Niblack thresholding from

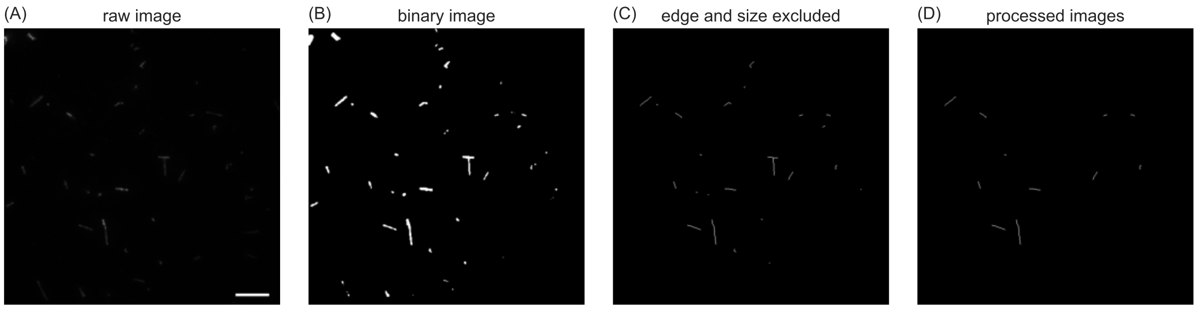

Fig. S1: **Processing steps of microtubule images.** (A) Raw image. Scale bar denotes 10 µm. (B) Images processed after computing a Niblack threshold and using Otsu thresholding on the Niblack threshold array. (C) Putative MTs skeletonized after removing objects too close to the image border or too small. (D) Removal of any MTs that cross over each other to get the final MTs used for analysis.

being picked up as signal. This closing was performed using a 3 pixel x 3 pixel square array, suggesting that disconnected microtubules needed to be within $3\sqrt{2}$ pixels of each other at their ends to be connected again. From here, we removed any microtubules that were too close to the edge of the image as they may extend outside of the camera field of view and removed any objects that were fewer than 10 pixels in area as we considered them too small to know with enough certainty whether they were microtubules or small blemishes in the image. Putative microtubules underwent a morphological thinning so that they were converted to one-pixel wide lines along which we could compute their lengths. The result of the edge and size exclusion and skeletonizing are shown in Fig. S1(C).

As a final step before measuring the lengths, we removed any microtubules that seem to cross over. This was performed by removing objects where two line segments along the same microtubule strand formed angles of at least 75°, leaving behind a processed image such as Fig. S1(D). From here, we used any remaining microtubules and measured their lengths and compiled them. Fig. S2 shows empirical cumulative distribution functions of these microtubules from the five MT polymerization assays performed over the course of the work presented here. $n$ denotes the number of microtubules that were extracted from the image processing and used in the distributions for each replicate. Here, we see that for most of the work performed the MTs had lengths between $1-3$ µm with median lengths between $1.5-2.1$ µm.

#### S3 Image processing: global drift correction

For computational efficiency in later image processing steps, photobleached images are cropped to contain only the region where the collective filament network is present. We first find the center of the contracting network for the image immediately preceding photobleaching. To do so, the image is smoothed with a Gaussian blur and thresholded with the Yen thresholding method [7]. After removing objects that are at the image edge or small objects, the largest segmented object is taken. The properties of this object are then taken, included a pixel-weighted centroid and its major axis length.

For cropping the first photobleached image, we start by cropping a window in the image using the pixel-weighted centroid as the image center and 130% of the major axis length as the length of the window. This buffer to the window cropping typically ensures that contracting networks that are drifting can still be easily tracked and cropped. To more efficiently crop the image, we then take this cropped window and use a heavy Gaussian blur ($\sigma = 30$ pixels) and subtract this from the cropped photobleached image. We then normalize the image and use Otsu thresholding [5] to identify putative fluorescent unit cells. We roughly identify the unit cells by removing those that are close to the edge of the image as well as objects that are smaller than 36 µm$^2$, which would be far smaller than a unit cell. Unit cells are further cleaned up by filling in any small holes in the unit cell with a morphological closing before taking the pixel-weighted centroid using all of the unit cells together to get a rough position of the network center. This process is then repeated on the next photobleached frame using the new centroid for the image center and the original window length over the desired number of photobleach frames. These cropped images are then used for further, more careful processing of the unit cells.

#### S4 Quantifying microtubule unit cell dynamics

In this work, we sought to characterize the bulk redistribution of microtubules through local deformations and translations within the contracting network. To develop a processing method that would allow us to quantify the advective and diffusive components of the network, we first set out to determine whether the microtubule number is conserved in the system. A part of this determination, which relies heavily upon the fluorescence signal, depends upon whether the imaging system

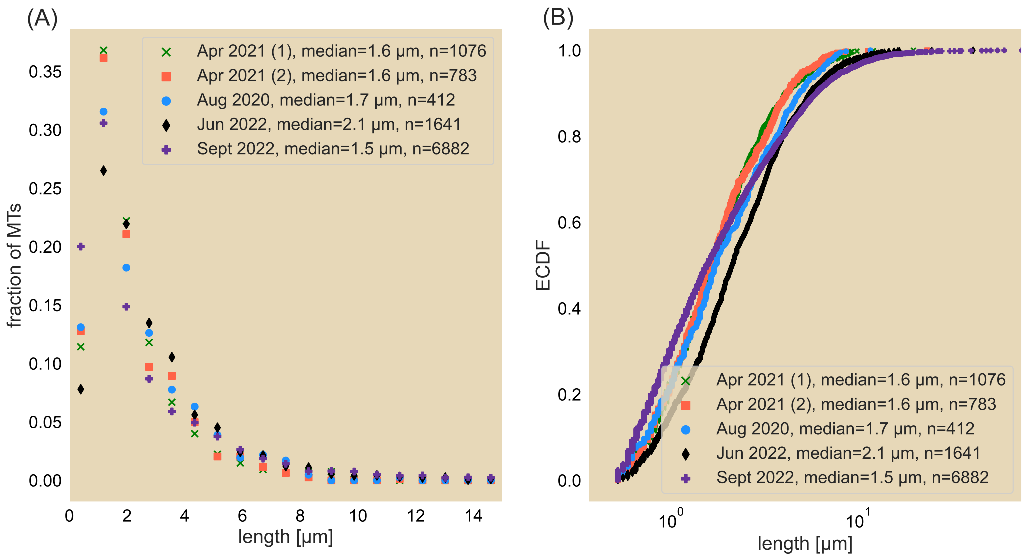

Fig. S2: **Distributions of microtubule length from microtubules stabilized from polymerization preparations for experiments used in this manuscript.** Microtubules were prepared five times over the course of the work presented here, thus shown as five different datasets. Left plot shows the histogrammed length distribution as a linear x-scale of length while the right plot shows the same data as an empirical cumulative distribution function (ECDF) as a logarithmic x-scale. The two polymerization preparations performed in April 2021 were performed separately by two of the authors of this manuscript on the same day. $n$ denotes the number of microtubules whose lengths were obtained in the distributions.

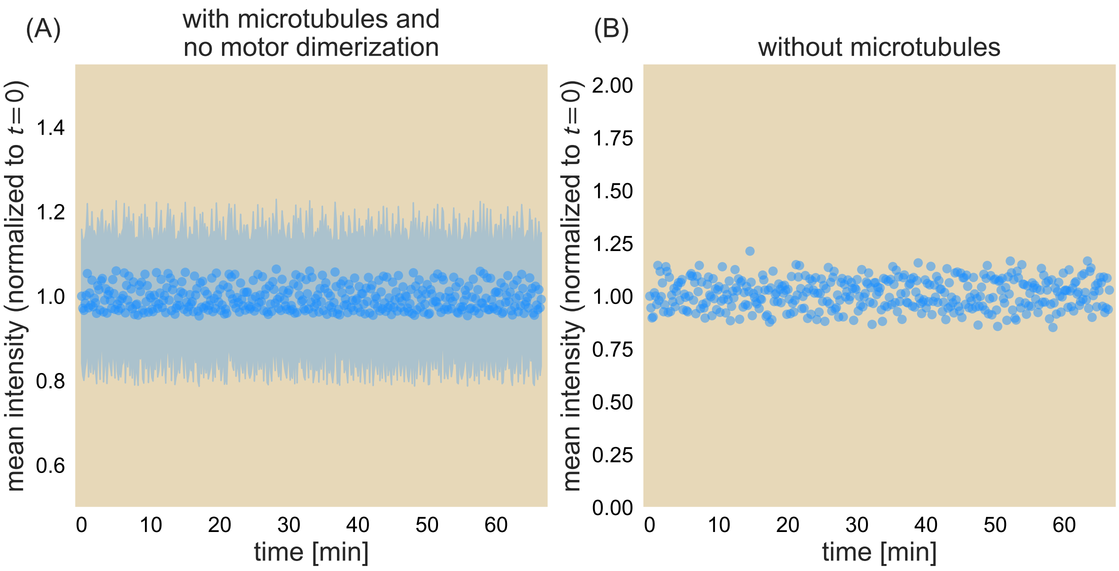

Fig. S3: **Image intensity of the microtubule field as a function of time.** (A) Mean intensity of the microtubule field normalized against that of the first image. Blue shaded region represents one standard deviation in the mean intensity (normalized by the same initial mean value). (B) Mean intensity of the same fluorescence channel in the absence of microtubules.

also affects the signal over time through passive photobleaching. We seek to assess these factors in SI Secs. S4.1 and S4.2 below.

#### S4.1 Imaging system negligibly photobleaches microtubules.

One concern in analyzing microtubule fluorescence over time is whether the optical system decreases its signal due to secondary photobleaching effects from the projector, which is used to illuminate the field of view for imaging purposes and perform the iLid-micro light stimulation. To investigate this, we imaged the microtubule field without activating the iLid-micro dimerization using the same exposure times (200 ms) and different imaging frequencies depending upon the speed at which contraction takes place, between 3 seconds and 10 seconds per frame. We then examined the mean image intensity and standard deviation of the pixel intensity as a function of time.

Fig. S3(A) illustrates the effects of the projector on the microtubule field. The mean intensity of the field of view, as normalized against the mean intensity at $t = 0$ seconds, indicates that the fluorescence field fluctuates within a few percent but does not appear to decrease over an hour. These fluctuations are likely due to fluctuations from the image acquisition set-up itself, as Fig. S3(B) shows the normalized mean intensity of the microtubule fluorescence channel but in the absence of microtubules. Here, we see that that the integrated intensity fluctuates over the short term but does not appear to exhibit a global decrease, further supporting that the small fluctuations in fluorescence intensity in successive imaging stages comes from the imaging system. Nevertheless, we conclude that the fluorescence intensity is well preserved over the course of experiments and does not require corrections during image processing.

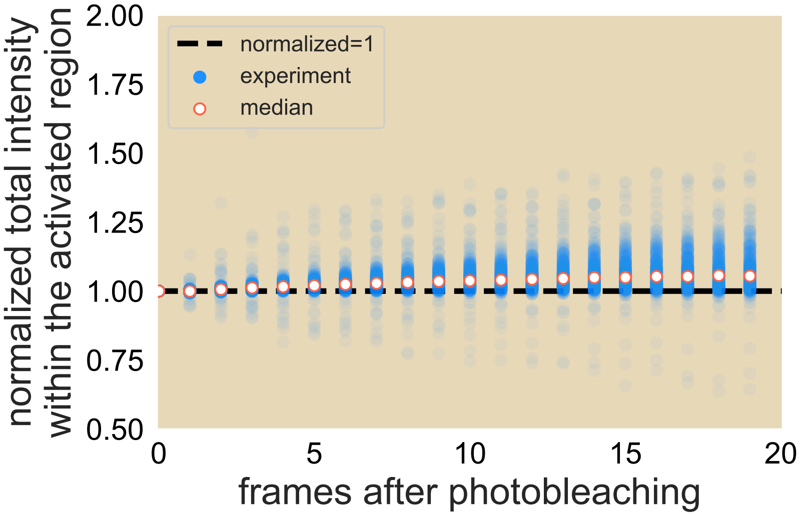

Fig. S4: **Integrated intensity of the photobleached contracting network over image frames.** White dots denote the median value across all experiments.

#### S4.2 Net flux of microtubules goes into the imaging plane.

As the projector does not passively photobleach the microtubule channel (SI Sec. S4.1), we next ask whether there is a loss of microtubules during the contraction process. Microtubules may disconnect from the contracting network and diffuse away. For example, as we use an epifluorescent imaging set-up, if microtubules are lost from the network by moving out of the plane of imaging, they will project a low, more diffuse signal onto the image. In contrast, microtubules that move into the plane of focus will exhibit a higher signal. Similarly, microtubules lost at the periphery in the image plane will lead to a reduced integrated intensity across the entire network.

To determine the effects of flux across the focal plane, we measure the integrated fluorescence signal of the contracting network after photobleaching. This is done by integrating the fluorescence signal of the activated network as it contracts away from the unilluminated reservoir. We then examine the normalized integrated signal over time.

Fig. S4 shows that the integrated intensity of the contracting network increases in time. Over 20 imaging frames, the fluorescence increases by about 5%, suggesting a roughly 0.25% increase between frames. While some of the observed increase in intensity can be accounted for by those datasets where the contracting network is not fully disconnected from the unilluminated reservoir and thus introducing more microtubules at the periphery, we suspect that the majority of this increase comes from an increase of microtubules that are entering the imaging plane. This observation makes sense as we expect a growing concentration of microtubules entering the imaging plane due to network contraction. Had we accounted for this increase in intensity over successive frames, our results would at best have led to a greater area of the unit cells than the ones we computed, which would produce greater effective diffusion constants. Even so, we argue that the roughly quarter of a percent increase between frames is relatively minor and conclude that the total microtubule count remains roughly constant over the course of the experiment.

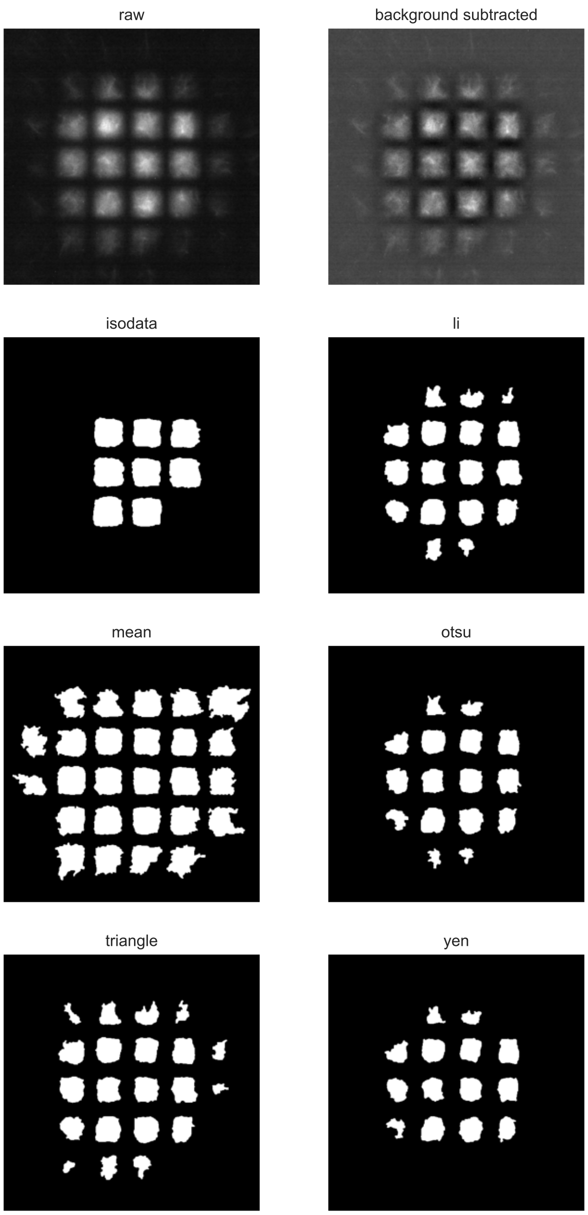

Fig. S5: **Various thresholding schemes of fluorescent squares.** Top two images correspond to the raw (left) and background-subtracted (right) images. The thresholding schemes used, in order, were isodata, Li, mean, Otsu, triangle, and Yen thresholding methods. Due to the under-representation of unit cell fluorescent signal for all the other methods, we opt for the mean thresholding scheme to identify unit cells.

##### S4.3 Number conservation of unit cells

Due to the negligible photobleaching effects of the projector on the network and the small influx of microtubules in the imaging plane, we make the assumption that the total number of microtubules for the entire network is conserved. We further assume that at the local level, the number of fluorescent microtubules that compose a unit cell is also conserved. As a result, we choose to identify and track unit cells in time by conserving their integrated fluorescence intensity. Fluorescent unit cells of a photobleached microtubule network are thus segmented in the cropped image sets where the microtubules outside of the activation region are neglected. For each image, we identify the unit cells by first enhancing the contrast between the fluorescence signal of the unit cells and the background through the subtraction of a heavily Gaussian blurred form of the image ($\sigma = 20$ pixels) and subtracting off this blur from the original image. Pixel values are then normalized across the image to fall between 0 and 1.

In order to identify each fluorescent square, we tested a variety of thresholding schemes using the Sci-kit Image package for Python. In summary, the following thresholding schemes are:

- Isodata - identifies those threshold values where, when each pixel is grouped according to whether it lies above or below the threshold value, the threshold value is the average of the two binned groups.
- Li - iteratively computes the cross-entropy between the image and a binary image with a different thresholding value. The returned threshold value is that which reduces the cross entropy [4].
- Mean - computes the mean pixel value across the image [8].
- Otsu - finds the threshold that minimizes the sum of the variances of the background and foreground [5].
- Triangle - computes a line from the peak in the histogram to the last histogram bin (if the peak is shifted to the left of the histogram). A second perpendicular line is drawn from this line toward the first histogram bin it touches. The corresponding x-value gives the threshold [6].
- Yen - computes the minimum cross-entropy between the image and thresholded binary image while accounting for the bit depth of the image [7].

Furthermore, we seek the method that best identifies the unit cells not only in the bulk that will appear as squares but also those that lie along the periphery that may not appear as complete squares after the photobleaching is applied but are nevertheless part of the network.

Fig S5 shows these various thresholding schemes performed on the background-subtracted image (top right) with comparisons to the original raw image (top left). We see that while the isodata thresholding approach misses many of the fluorescent squares, other thresholding schemes reasonably render the threshold of the squares. We notice that the Li, Otsu, triangle and Yen thresholding schemes miss unit cells on the periphery of the network, especially if they are not squares as in those found toward the center of the microtubule network. To keep track of their dynamics, we elect to use the mean thresholding algorithm, which from visually comparing the threshold to the raw data better represents the unit cells, including those unit cells on the network periphery. After the thresholding is applied, the segmented image is cleaned up by removing segmented objects that are too small (less than a third of the area of a unit cell immediately after photobleaching) and objects that are larger than the area of a unit cell. A morphological closing is performed where any holes smaller than 3 pixels $\times$ 3 pixels within a fluorescent unit cell is closed. These small holes may arise from a local minimum in signal that falls in the background regime during thresholding. With the segmented images from the first frame, the centroid position, area, and total fluorescence of each unit cell are computed. For total fluorescence, we compute the pixel intensity by taking the raw image signal and subtracting the average background signal inherent to the camera. Fig S6(A) provides a schematic of the resultant thresholding to initially identify unit cells.

As schematized in Fig S6(B) and (C), images of subsequent time points are processed with the intention of preserving the integrated fluorescence of each unit cell, which corresponds with our argument that fluorescent microtubules are conserved for each unit cell. We first subtract the background signal and segment unit cells with the same threshold value as for the $t = 0$ timepoint. However, as unit cells begin to deform due to their fluorescent microtubules dispersing, the integrated fluorescence signal of the newly segmented unit cells will differ from that of the first time point, which translates to a different number of fluorescent microtubules. As a result, for images after the first frame immediately succeeding photobleaching, we expand or reduce the segmentation of unit cells by adding or subtracting pixels around their boundaries until we obtain the same total fluorescence as the $t = 0$ timepoint. Fig. S6(C) elaborates on the scheme for correcting to obtain the same integrated intensity as in the initial time frame to within a specified tolerance. To do this, each unit cell is then paired with itself from the previous time step by determining nearest centroids. Due to the minimal reduction in fluorescence intensity from the projector during imaging as discussed in Sec S4.1, we compare the total fluorescence intensity of the segmented unit cell in the frame of interest to that of the same unit cell from the first frame. 1 layer of pixel beyond (within) the boundary of the unit cell are histogrammed and Otsu thresholded to distinguish microtubule regions to background regions. The pixels that make up the foreground (background) according to the thresholding are then added (subtracted) until the integrated fluorescence falls within 0.01 tolerance of the original fluorescence intensity.

To understand how the choice of relative tolerance in the integrated fluorescence affects that computed effective diffusion constant $D_{\text{eff}}$, we performed the unit cell segmentation and tracking under different tolerance levels. Fig S7 shows that while a tolerance below 0.015 leads to a constant effective diffusion constant, increasing the tolerance above this point leads to a monotonic decrease in the $D_{\text{eff}}$. This suggests that the area trajectories of unit cells can be highly sensitive to the tolerance given to the microtubule preservation count. This control also indicates that the area trajectories of unit cells are not markedly different below the 0.015 tolerance and yields robust measurements of the area trajectories and by extension fits of the effective diffusion constants.

Unit cell centroids, areas, and fluorescence intensities are then computed in addition to the pixel-weighted center of the entire contracting network after this intensity-adjusted processing for all of the unit cells. Image processing of a unit cell terminates when it is found to overlap with another unit cell during the fluorescence intensity correction scheme as this indicates that the unit cells have begun to merge and by the next time point thus microtubules from one unit cell can no longer be distinguished from those of the other.

### S5 Data analysis

#### S5.1 Contraction rate computation

In the main text, we use the centroids of fluorescent unit cells obtained in SI Sec. S4.3 to demonstrate that contraction speed of the microtubule network scales linearly with distance from the network center. We first obtain the speed that each unit cell centroid is moving toward the center as a function of time. For each unit cell, we observe a linear relation between the centroid distance

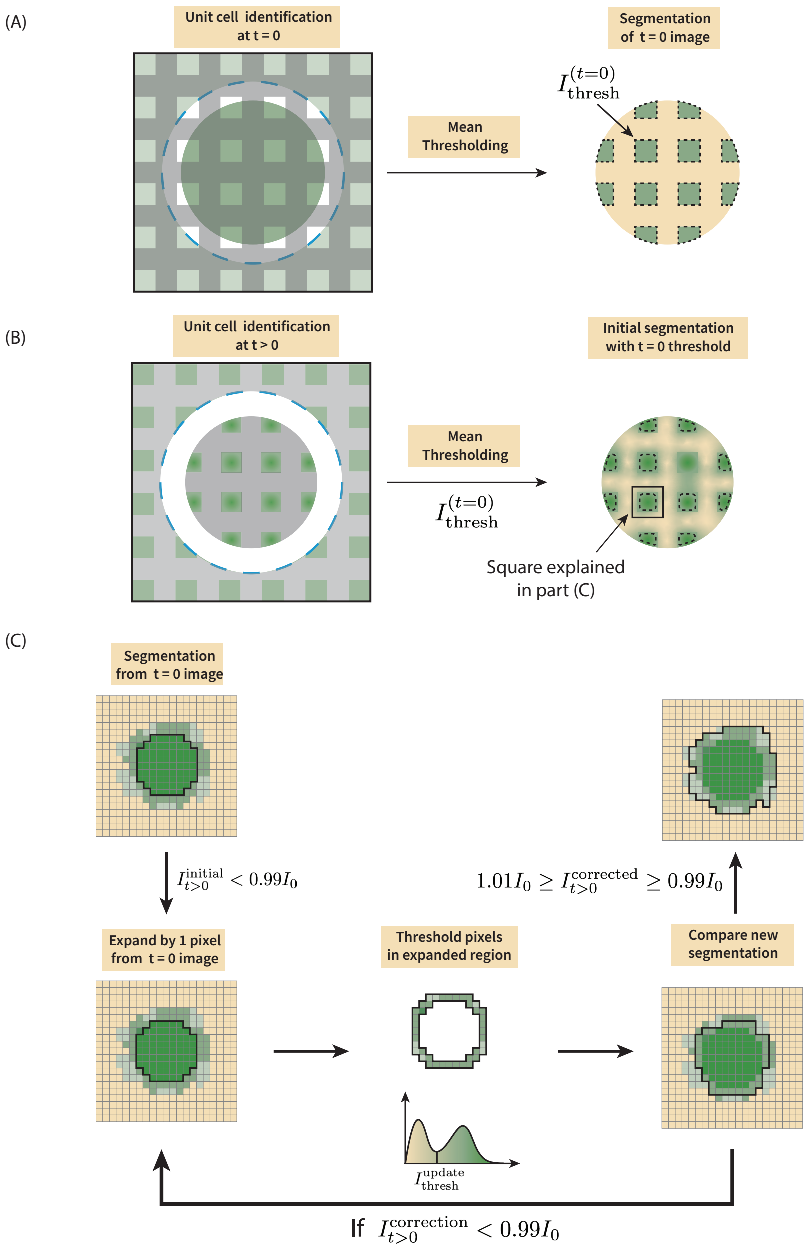

Fig. S6: **Unit cell segmentation correction scheme.** (A) Unit cells in the first image after photobleaching are segmented using mean thresholding to obtain an initial threshold value $I^{(t=0)}_{\mathrm{thresh}}$. Dashed blue circle denotes the extent of the projected light within which motors dimerize, causing the network to couple and contract (green circle). (B) Unit cells of later frames are initially segmented using $I^{(t=0)}_{\mathrm{thresh}}$. (C) The integrated intensity of each unit cell after the initial segmentation $I^{\mathrm{initial}}_{t>0}$ is compared against that for the $t = 0$ case, $I_0$. In instances where $I_{t>0} < 0.99 I_0$, the pixels in a single layer beyond the segmentation boundary are histogrammed and thresholded to distinguish pixels containing microtubules with those regions that make up the background. These pixels with signal are then added, the integrated intensity is recomputed and compared again to $I_0$. The process is repeated until the integrated intensity falls within 1% of $I_0$.

from the network center and time after photobleaching of the form

$$r = v_c t + r_0, \tag{S1}$$

where $r$ is the unit cell centroid distance from the network center, $v_c$ is the speed of the unit cell (which will take to be positive here but directed toward the origin), $t$ is the time since photobleaching, and $r_0$ is the initial centroid distance from the network center immediately after photobleaching.

Based on the extracted contraction speed and distances for all of the unit cells for a given motor type, we noted a linear relation between radius $r$ and centroid speed $v_c$ of the form

$$v_c = \alpha r + v_0, \tag{S2}$$

where $\alpha$ is the contraction rate in units of inverse time and $v_0$ is the contraction speed at the network center. To this end, we aim to compute $\alpha$ and $v_0$. Although we expect the speed at the network center to be 0, we relax this assumption for our analysis. To more carefully compute the rate of contraction of the network and determine the range of credibility of the computed rate, we use a Bayesian approach. Specifically, we compute the probability of $\alpha$ and $v_0$ given the contraction speed and distance of each unit cell from the network center $(r_0, v_c)_i$, $P(\alpha, v_0| \{(r_0, v_c)_i\})$, where $i$ denotes each unit cell. Here, we use the centroid distance immediately after photobleaching but found that another criterion such as the median of the centroid distance over the course of the time window analyzed does not dramatically affect the results due to the relatively small travel ($\frac{\Delta r}{r_0} < 10\%$ for $\Delta r$ the distance traveled over the entire time course) the unit cells undergo.

We note from Bayes' Theorem that

$$P[\alpha, v_0| \{(r_0, v_c)_i\}] = \frac{P[\{(r_0, v_c)_i\} |\alpha, v_0] \, P(\alpha, v_0)}{P[\{(r_0, v_c)_i\}]},$$

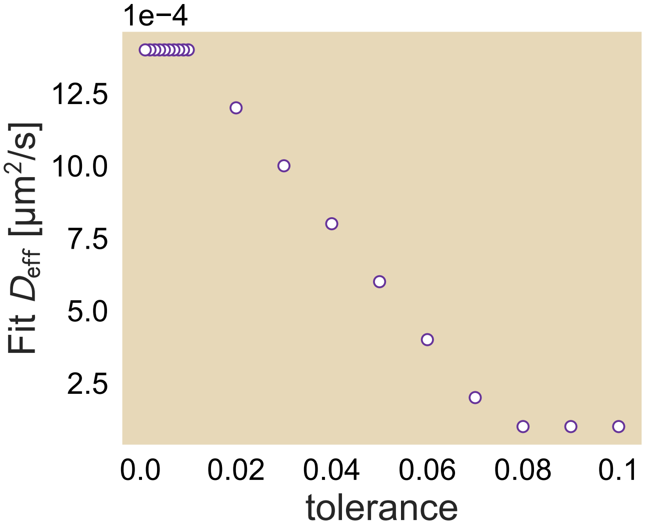

Fig. S7: **Effective diffusion constant fits against various tolerances in the relative unit cell fluorescence.** The tolerance is the fractional difference in fluorescence intensity between the unit cell in the first frame and the unit cell at a later time point. Dataset used on Ncd236 at saturated ATP concentration (1.4 µM).

$$
\begin{aligned}
&= \frac{\prod_i P\left[(r_0, v_c)_i \,|\alpha, v_0\right]}{\prod_i P\left[(r_0, v_c)_i\right]} P\left(\alpha, v_0\right), \\
&\propto \prod_i P\left[(r_0, v_c)_i \,|\alpha, v_0\right] P\left(\alpha, v_0\right),
\end{aligned}
\tag{S3}
$$

where we drop the denominator on the right-hand side as it does not involve the parameters we want to find, thus making the two sides proportional to each other. Here, $P\left[(r_0, v_c)_i \,|\alpha, v_0\right]$ is the likelihood distribution of getting the $(r_0, v_c)_i$ that we did given $\alpha$ and $v_0$ while $P\left(\alpha, v_0\right)$ is the prior distribution of our two parameters.

We expect that our priors on $\alpha$ and $v_0$ are independent of each other, so we can break up the probability function into a product of two functions

$$
P\left(\alpha, v_0\right) = p\left(\alpha\right) p\left(v_0\right). \tag{S4}
$$

Meanwhile, we can rearrange each likelihood function as a product of two probabilities. The probability of getting $(r_0, v_c)_i$ given our parameters is also the probability of getting $v_{c,i}$ given our parameters and $r_{0,i}$ times the probability of getting $r_{0,i}$, or

$$
\begin{aligned}
P\left((r_0, v_c)_i \,|\alpha, v_0\right) &= P\left(v_{c,i}|\alpha, v_0, r_{0,i}\right) P\left(r_{0,i}\right), \\
&\propto P\left(v_{c,i}|\alpha, v_0, r_{0,i}\right),
\end{aligned}
\tag{S5}
$$

where we change to a proportionality again as $P\left(r_{0,i}\right)$ is independent of our parameters. Here, we expect that our contraction speed for a given unit cell $v_{c,i}$ comes from a Normal distribution where the mean value is $\alpha\, r_{0,i} + v_0$ and standard deviation $\sigma$. This means that we will also need a prior on $\sigma$. This means that our distribution really takes the form of

$$
P\left(\alpha, v_0, \sigma | \left\{(r_0, v_c)_i\right\}\right) \propto P\left(\alpha\right) P\left(v_0\right) P\left(\sigma\right) \prod_i P\left(v_{c,i}|\alpha, v_0, \sigma, r_{0,i}\right). \tag{S6}
$$

As a result, we say that our likelihood takes the form

$$
v_{c,i} \backsim \text{Normal}\left(\alpha r_{0,i} + v_0, \sigma^2\right). \tag{S7}
$$

We then defined our priors to be that $\alpha$ is drawn from the half-normal distribution where $\alpha > 0$ as we are working with speeds of contraction, $\sigma$ is also drawn from a half-normal distribution and enforced to be positive, and $v_0$ is drawn from a normal distribution about $v = 0$. We make the offset a normal rather than a half-normal distribution as there may be a value of $r > 0$ for which the contraction stops, which for a positive slope would mean a negative speed at $r = 0$. Put together, we have the following priors

$$
\alpha \backsim \text{Half-Normal}\left(0, 1\right), \tag{S8}
$$

$$
\sigma \backsim \text{Half-Normal}\left(0, 1\right), \tag{S9}
$$

$$
v_0 \backsim \text{Normal}\left(0, 1\right). \tag{S10}
$$

We sampled the joint distribution of $(\alpha, v_0, \sigma)$ by Hamiltonian Markov chain Monte Carlo using the Stan probabilistic program [9]. From each $(\alpha, v_0)$ that is sampled we compute the mean value $\mu = \alpha\, r + v_0$ for $0 \leq r \leq R$ where $R$ is the distance of the farthest centroid from the network center and report the median and 95% credible region in Fig. 2 and 4-6 of the main text.

##### S5.2 Computing the best fit effective diffusion constant

In the main manuscript, we use an advection-diffusion model to compute an effective diffusion constant to quantify the difference in area between the experimental normalized area trajectories and the pure contraction bound (signifying no diffusion). To do so, we used the finite element method (FEM) on individual unit cells of initially uniform concentration subject to the advection-diffusion equation as described in Eq. 3 of the main manuscript. We then processed the simulated concentration field data with a similar integrated particle count method as described in SI Sec. S4.3 in order to compute the area of the unit cells in time. This analysis gives rise to a family of normalized area trajectories for a fixed contraction rate and variable diffusion constant. In order to compute the effective diffusion constant from, say, the median normalized area trajectories from a given set of experimental conditions, we take the simulated area trajectory for one of the diffusion constants and the area trajectory of the experimentally-obtained contraction rate and compute the sum of the square of the difference between the two trajectories across time. For each of the quartiles, the effective diffusion constant is computed as the one whose area trajectory minimizes the sum of the differences squared.

#### S6 Experimental variation of contraction speed and normalized area trajectories

In the main manuscript, we computed the contraction rate using all of the replicates of a given set of experimental conditions. However, to exhibit experimental variation between replicates, we present in Fig. S8 the contraction speed and normalized area data for all of the replicates involving Ncd236 at 1400 µM ATP and 1.5 mg/mL pluronic. Note that the line in the contraction speed is the same as shown in Fig. 2(D) of the main manuscript where the contraction rate $\alpha = 0.002$ sec$^{-1}$ for comparison of how each replicate compares to the computed line. This contraction is also used for the pure contraction bound shown on the normalized area data. The time noted at the top of each contraction speed plot marks the time into the experiment that the photobleaching was performed, with the plots organized in order of ascending time into the experiment of photobleaching.

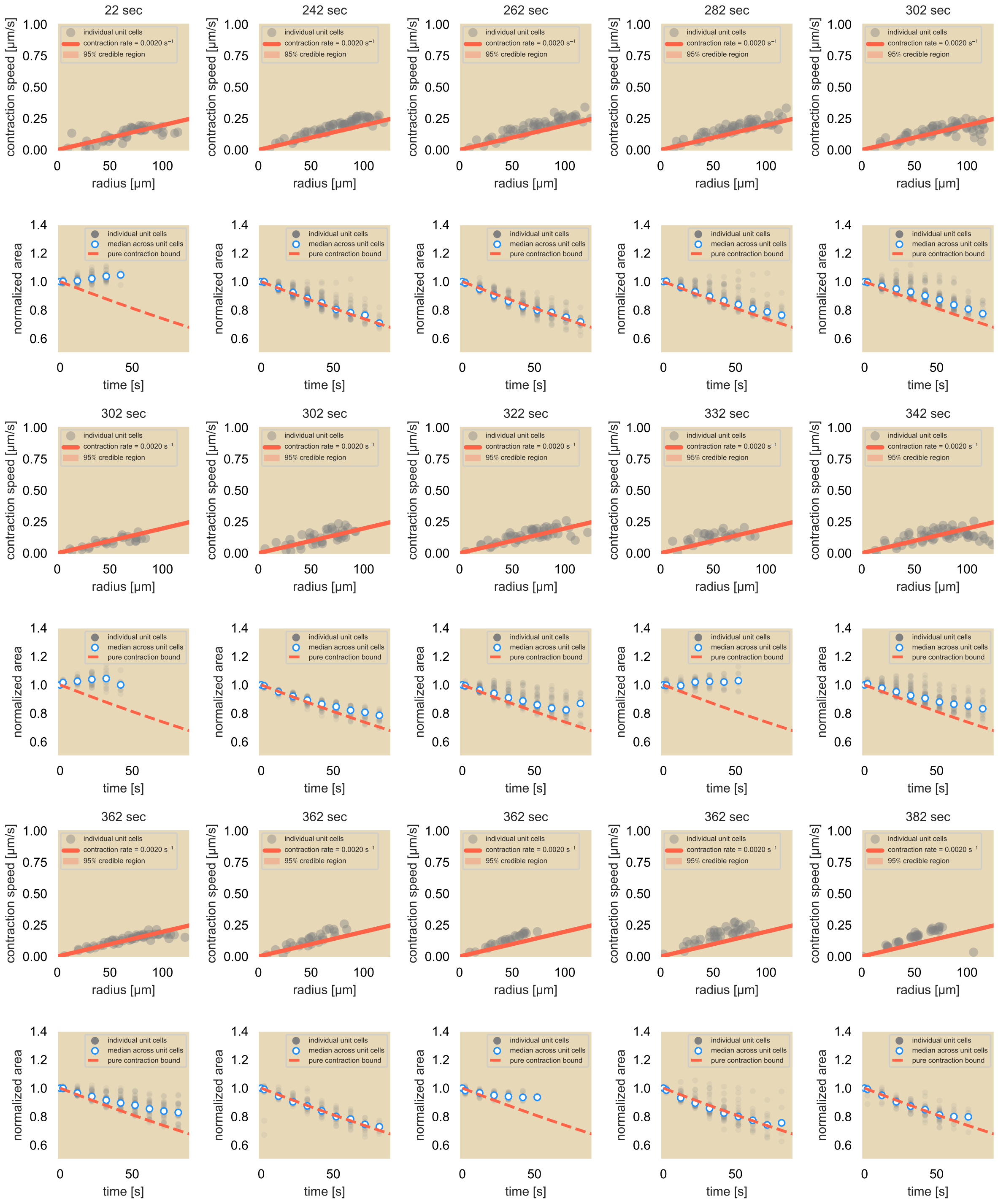
22 sec
242 sec
262 sec
282 sec
302 sec
302 sec
302 sec
322 sec
332 sec
342 sec
362 sec
362 sec
362 sec
362 sec
382 sec
individual unit cells
contraction rate = 0.0020 s⁻¹
95% credible region
contraction speed [µm/s]
radius [µm]
median across unit cells
pure contraction bound
normalized area
time [s]

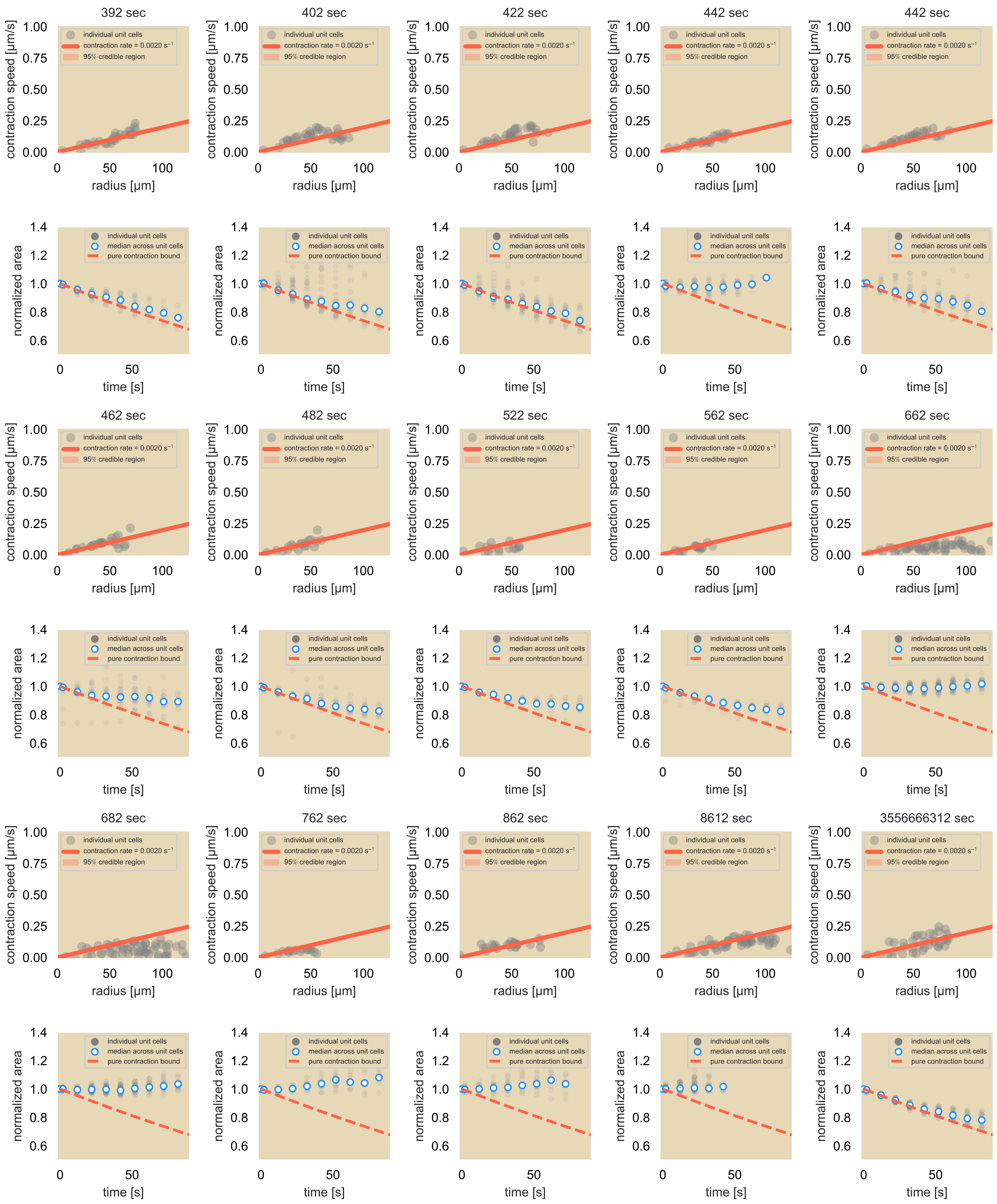

Fig. S8: **Contraction speed (odd rows) and normalized area trajectory (even rows) of each experimental replicate using 1.5 mg/mL pluronic, 1400 µM ATP, and Ncd236.** The lines in the plots of contractions speed data and in the plots of the area trajectory are the same as in Fig. 2(D) and 2(F), respectively, of the main manuscript. The time at the top of each contraction speed plot marks the time into the experiment that the photobleaching was performed.

#### S7 Deformation of a square due solely to contraction

In the main text, we observed that each fluorescent unit cell on average conserves its area while its center of mass moves toward the network center with speed that is linearly dependent on the distance from the center. We compute the expected area of each unit cell had the network elastically contracted due solely to the observed global contraction. We define the contraction velocity field $\mathbf{v}(x, y)$ as

$$\mathbf{v}(x, y) \equiv -\alpha \left(x\hat{x} + y\hat{y}\right), \tag{S11}$$

where $\alpha$ is the contraction rate as computed in SI Sec. S5.1 and reported in the main manuscript. This means that after a time interval $\Delta t$ a point $(x, y)$ subject to this advective flow will be displaced in the x- and y- directions according to

$$\begin{aligned} \Delta X &= v_x \Delta t = -\alpha x \Delta t, \\ \Delta Y &= v_y \Delta t = -\alpha y \Delta t, \end{aligned} \tag{S12}$$

so the point at the later time $(x', y')$ relates to its earlier time point by

$$\begin{aligned} x' &= x + \Delta X = x\left(1 - \alpha\Delta t\right) \\ y' &= y + \Delta Y = y\left(1 - \alpha\Delta t\right). \end{aligned} \tag{S13}$$

Suppose we looked at the four corners of a unit cell, labeled as A, B, C, D as depicted in Fig. S9. If we assign their coordinates as

$$\begin{aligned} \mathrm{A} &\to \left(x_\mathrm{A}, y_\mathrm{A}\right), \\ \mathrm{B} &\to \left(x_\mathrm{B}, y_\mathrm{B}\right), \\ \mathrm{C} &\to \left(x_\mathrm{C}, y_\mathrm{C}\right), \\ \mathrm{D} &\to \left(x_\mathrm{D}, y_\mathrm{D}\right). \end{aligned} \tag{S14}$$

Under a rectangular geometry, we can choose two vertices diagonally across from each other on the rectangle and write their x- and y- coordinates with the coordinates of the other two diagonal vertices, so with a choice of using coordinates from A and D, the coordinates for B and C become

$$\begin{aligned} \mathrm{A} &\to \left(x_\mathrm{A}, y_\mathrm{A}\right), \\ \mathrm{B} &\to \left(x_\mathrm{D}, y_\mathrm{A}\right), \\ \mathrm{C} &\to \left(x_\mathrm{A}, y_\mathrm{D}\right), \\ \mathrm{D} &\to \left(x_\mathrm{D}, y_\mathrm{D}\right). \end{aligned} \tag{S15}$$

Under the deformation mapping, their new coordinates, labeled as A′, B′, C′, and D′ get mapped on as

$$\begin{aligned} \mathrm{A}' &\to \left[x_\mathrm{A}\left(1 - \alpha\Delta t\right), y_\mathrm{A}\left(1 - \alpha\Delta t\right)\right], \\ \mathrm{B}' &\to \left[x_\mathrm{D}\left(1 - \alpha\Delta t\right), y_\mathrm{A}\left(1 - \alpha\Delta t\right)\right], \\ \mathrm{C}' &\to \left[x_\mathrm{A}\left(1 - \alpha\Delta t\right), y_\mathrm{D}\left(1 - \alpha\Delta t\right)\right], \\ \mathrm{D}' &\to \left[x_\mathrm{D}\left(1 - \alpha\Delta t\right), y_\mathrm{D}\left(1 - \alpha\Delta t\right)\right]. \end{aligned} \tag{S16}$$

Eqs. S16 tells us that under this particular velocity field, any two points that are horizontally or vertically aligned will maintain the same horizontal or vertical alignment, respectively, even at later times. Thus, a square will preserve its shape in time.

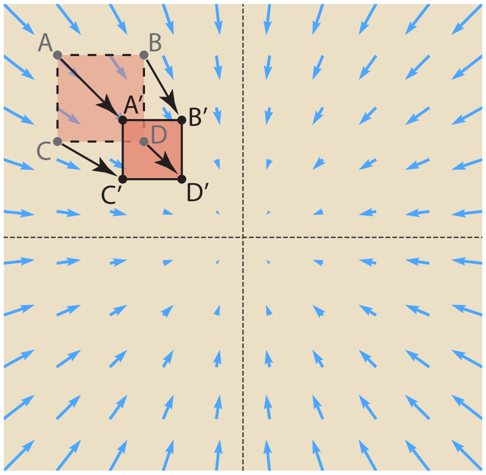

Fig. S9: **Schematic of unit cell contraction due purely to the advective velocity field.** An advective velocity field scales linearly with distance from the origin while pointing radially inward and is shown in blue. The points at the corners of the square (A, B, C, D) are mapped after some time $\Delta t$ to the points (A′, B′, C′, D′).

We next examine what happens to the area of a unit cell had the only effect been the global contraction. In this case, we can compare the area of the square before and after the deformation. To compute the area swept out by (A,B,C,D), we multiply the line segment between B and D, $L_{\mathrm{BD}}$ with the line segment between C and D, $L_{\mathrm{CD}}$:

$$\begin{aligned}\sigma_{(\mathrm{A,B,C,D})} &= L_{\mathrm{BD}} \times L_{\mathrm{CD}}, \\ &= \left[\sqrt{(x_{\mathrm{B}} - x_{\mathrm{D}})^2 + (y_{\mathrm{B}} - y_{\mathrm{D}})^2}\right] \times \left[\sqrt{(x_{\mathrm{D}} - x_{\mathrm{C}})^2 + (y_{\mathrm{D}} - y_{\mathrm{C}})^2}\right], \\ &= (y_{\mathrm{A}} - y_{\mathrm{D}}) \times (x_{\mathrm{D}} - x_{\mathrm{A}})\,. \end{aligned} \tag{S17}$$

As noted from Eq. S15, we used the fact that $x_{\mathrm{B}} = x_{\mathrm{D}}$, $y_{\mathrm{B}} = y_{\mathrm{A}}$, $x_{\mathrm{C}} = x_{\mathrm{A}}$, and $y_{\mathrm{C}} = y_{\mathrm{D}}$ to simplify the equation down in terms of two coordinates. In comparison, the area of the deformed unit cell swept out by (A′, B′, C′, D′) takes the form

$$\begin{aligned}\sigma_{(\mathrm{A',B',C',D'})} &= L_{\mathrm{B'D}} \times L_{\mathrm{C'D'}}, \\ &= \left[\sqrt{(x_{\mathrm{B'}} - x_{\mathrm{D'}})^2 + (y_{\mathrm{B'}} - y_{\mathrm{D'}})^2}\right] \times \left[\sqrt{(x_{\mathrm{D'}} - x_{\mathrm{C'}})^2 + (y_{\mathrm{D'}} - y_{\mathrm{C'}})^2}\right], \\ &= (y_{\mathrm{A'}} - y_{\mathrm{D'}}) \times (x_{\mathrm{D'}} - x_{\mathrm{A'}})\,, \\ &= [y_{\mathrm{A}}\,(1 - \alpha\Delta t) - y_{\mathrm{D}}\,(1 - \alpha\Delta t)] \times [x_{\mathrm{D}}\,(1 - \alpha\Delta t) - x_{\mathrm{A}}\,(1 - \alpha\Delta t)]\,, \\ &= (y_{\mathrm{A}} - y_{\mathrm{D}})\,(1 - \alpha\Delta t) \times (x_{\mathrm{D}} - x_{\mathrm{A}})\,(1 - \alpha\Delta t)\,, \\ &= (y_{\mathrm{A}} - y_{\mathrm{D}}) \times (x_{\mathrm{D}} - x_{\mathrm{A}})\,(1 - \alpha\Delta t)^2\,, \\ &= \sigma_{(\mathrm{A,B,C,D})}\,(1 - \alpha\Delta t)^2\,. \end{aligned} \tag{S18}$$

Thus we find that the area of the unit cell subject solely to the contraction would decrease by $(1 - \alpha\Delta t)^2$ after a time period $\Delta t$. This comes in contrast to the results that we found experimentally where the area of the fluorescent unit squares remains greater than the pure contraction bound during the contraction process suggesting a mechanism that disperses microtubules against the global contraction.

### S8 2D Linear Advection-Diffusion Model

In the work presented in the main manuscript, we argue for an advection-diffusion model to describe the redistribution of microtubules in the bulk of the contracting network. In this section, we explore Eq 3 as shown in the manuscript to examine whether the model could reasonably recapitulate the experimental observations as a tool for computing an effective diffusion constant for the diffusive-like spread of microtubules in the bulk. While the main manuscript uses the finite element method (FEM) to simulate the area change of a concentration of particles localized to a square, as is the case for microtubules of a unit cell, we first develop an intuition for this equation for a series of initial conditions and at steady state. To validate the FEM approach before applying it to the unit square case, we numerically and analytically solve these initial conditions and directly compare them. This theoretical analysis is meant to explore the filament concentration when subject to a linear contraction velocity profile. We note here that we later invoke the Sturm-Liouville Theorem, which we elaborate on in SI Sec S9.

In the 2D telescoping case, we assume that we are carrying out an aster assay experiment where we dimerize motors (and thus couple microtubules) in a circular region of radius $R$. We assume that the distributions of motors and microtubules are strictly radially dependent and thus have no angular dependence. Finally, we model the velocity profile of the microtubule movement by assuming radially inward advection of particles where those that lie further away from the origin move faster than those toward the center as given by

$$\mathbf{v} = -\alpha r \hat{r}. \tag{S19}$$

The advection-diffusion equation then takes the form

$$\begin{aligned}
\frac{\partial c}{\partial t} &= D\nabla^2 c - \nabla \cdot (\mathbf{v}c), \\
&= \frac{D}{r}\frac{\partial}{\partial r}\Big(r\frac{\partial c}{\partial r}\Big) + \alpha\frac{1}{r}\frac{\partial}{\partial r}(r^2 c), \\
&= D\frac{\partial^2 c}{\partial r^2} + \frac{D}{r}\frac{\partial c}{\partial r} + \alpha r\frac{\partial c}{\partial r} + 2\alpha c, \\
&= D\frac{\partial^2 c}{\partial r^2} + \Big(\frac{D}{r} + \alpha r\Big)\frac{\partial c}{\partial r} + 2\alpha c, \\
\frac{1}{D}\frac{\partial c}{\partial t} &= \frac{\partial^2 c}{\partial r^2} + \Big(\frac{1}{r} + \frac{\alpha r}{D}\Big)\frac{\partial c}{\partial r} + \frac{2\alpha c}{D}.
\end{aligned} \tag{S20}$$

We first follow the procedure of separation of variables $c(r,t) = \Phi(r)T(t)$ and determine that the time-dependent component takes on the familiar form of $e^{-Dk^2 t}$. This ansatz is then applied to Eq. S20 and we rewrite the spatial component of the concentration as

$$\begin{aligned}
-k^2\Phi &= \frac{\mathrm{d}^2\Phi}{\mathrm{d}r^2} + \Big(\frac{1}{r} + \frac{\alpha r}{D}\Big)\frac{\mathrm{d}\Phi}{\mathrm{d}r} + \frac{2\alpha\Phi}{D}, \\
0 &= r\frac{\mathrm{d}^2\Phi}{\mathrm{d}r^2} + \Big(1 + \frac{\alpha r^2}{D}\Big)\frac{\mathrm{d}\Phi}{\mathrm{d}r} + \Big(\frac{2\alpha}{D} + k^2\Big)r\Phi.
\end{aligned} \tag{S21}$$

We will define a new length scale $\lambda^2 \equiv \frac{D}{\alpha}$ as well as a change of variables $\rho \equiv \frac{r}{\lambda}$ and $\tilde{k} \equiv \lambda k$. In this case, Eq. S21 takes the altered form

$$0 = \rho \frac{\mathrm{d}^2\Phi}{\mathrm{d}\rho^2} + (1+\rho^2)\frac{\mathrm{d}\Phi}{\mathrm{d}\rho} + \left(2+\tilde{k}^2\right)\rho\Phi. \tag{S22}$$

By following a prescription on which we elaborate further in SI Sec S9, we obtain a weighting function that will help us compute the eigenfunctions using Eq. S36, namely,

$$m(\rho) = e^{\frac{\rho^2}{2}}. \tag{S23}$$

When we multiply Eq. S22 by the multiplicative function, we get

$$\begin{aligned} 0 &= \rho\, e^{\frac{\rho^2}{2}} \frac{\mathrm{d}^2\Phi}{\mathrm{d}\rho^2} + (1+\rho^2) e^{\frac{\rho^2}{2}} \frac{\mathrm{d}\Phi}{\mathrm{d}\rho} + \left(2+\tilde{k}^2\right)\rho\, e^{\frac{\rho^2}{2}}\Phi, \\ \frac{\mathrm{d}}{\mathrm{d}\rho}\left[\rho\, e^{\frac{\rho^2}{2}} \frac{\mathrm{d}\Phi}{\mathrm{d}\rho}\right] &+ 2\rho\, e^{\frac{\rho^2}{2}}\Phi = -\tilde{k}^2 \rho\, e^{\frac{\rho^2}{2}}\Phi. \end{aligned} \tag{S24}$$

Through the Sturm-Liouville Theorem as described in SI Sec S9, specifically Eq. S24, we find that the weighting function differs from the multiplicative function due to the inclusion of the prefactor $\rho$. In this case, the weighting function $w(\rho)$ as well as $p(\rho)$ and $q(\rho)$ are given as

$$w(\rho) = p(\rho) = q(\rho) = \rho\, e^{\frac{\rho^2}{2}}. \tag{S25}$$

Furthermore, we observe that the eigenvalues take the form $\tilde{k}^2$. Solutions of $\Phi$ from Eq. S24 are obtained from Wolfram Alpha and take the form

$$\begin{aligned} \Phi_{\mathrm{ss}}(\rho) &= c_{\mathrm{ss}}\, e^{-\frac{\rho^2}{2}}, \\ \Phi_{\mathrm{dyn}}(\rho) &= c_1\, e^{-\frac{\rho^2}{2}}\, {}_1F_1\Big(-\frac{\tilde{k}^2}{2}; 1; \frac{\rho^2}{2}\Big) + c_2\, G^{2,0}_{1,2}\left(\frac{\rho^2}{2}\middle|\begin{matrix}-\frac{\tilde{k}^2}{2}\\ 0,0\end{matrix}\right), \end{aligned} \tag{S26}$$

where $G^{m,n}_{p,q}\left(z\middle|\begin{smallmatrix}a_1,\ldots,a_p\\ b1,\ldots,b_q\end{smallmatrix}\right)$ is the Meijer G-function (we split up the eigenfunctions as dynamic and steady-state terms for now). We note here that the arguments of the Meijer G-function are such that the function diverges at the origin. As our system is defined as $0 \le r \le R$, we can say that $c_2 = 0$. Thus, our eigenfunctions are

$$\begin{aligned} \Phi_{\mathrm{ss}}(\rho) &= c_{\mathrm{ss}}\, e^{-\frac{\rho^2}{2}}, \\ \Phi_{\mathrm{dyn}}(\rho) &= c_1\, e^{-\frac{\rho^2}{2}}\, {}_1F_1\Big(-\frac{\tilde{k}^2}{2}; 1; \frac{\rho^2}{2}\Big), \end{aligned} \tag{S27}$$

where we note that in the case of $\tilde{k} = 0$, we go from the dynamic eigenfunction to the static eigenfunction.

#### S8.1 No-flux boundary condition

In the work presented here, there is no inflow or outflow of material at the boundary. Thus, we impose the boundary condition $\mathbf{J}\Big|_{r=R} = 0$. This means that

$$J_r\Big|_{r=R} = D\frac{\mathrm{d}\Phi}{\mathrm{d}r} - v(R)\Phi(R) = D\frac{\mathrm{d}\Phi}{\mathrm{d}r}\Big|_{r=R} + \alpha R\Phi(R) = 0. \tag{S28}$$

We then need to ensure that the boundary condition is satisfied for the dynamic eigenfunction. We start by taking the derivative of the eigenfunction:

$$
\begin{aligned}
\frac{\mathrm{d}\Phi}{\mathrm{d}r} &= -\frac{c_1\,\rho}{\lambda}\,e^{-\frac{\rho^2}{2}}\Big[\frac{\tilde{k}^2}{2}\,{}_1F_1\Big(1-\frac{\tilde{k}^2}{2};2;\frac{\rho^2}{2}\Big)+{}_1F_1\Big(-\frac{\tilde{k}^2}{2};1;\frac{\rho^2}{2}\Big)\Big],\\
\frac{\mathrm{d}\Phi}{\mathrm{d}r}\Big|_{r=R} &= -\frac{c_1\,\alpha R}{D}\,e^{-\frac{\alpha R^2}{2D}}\Big[\Big(\frac{Dk^2}{2\alpha}\Big){}_1F_1\Big(1-\frac{Dk^2}{2\alpha};2;\frac{\alpha R^2}{2D}\Big)+{}_1F_1\Big(-\frac{Dk^2}{2\alpha};1;\frac{\alpha R^2}{2D}\Big)\Big].
\end{aligned}
\tag{S29}
$$

so when applied to the boundary condition, we get

$$
\begin{aligned}
D\frac{\mathrm{d}\Phi}{\mathrm{d}r}\Big|_{r=R}+\alpha R\Phi(R) = &-c_1\,\alpha R\,e^{-\frac{\alpha R^2}{2D}}\Big(\frac{Dk^2}{2\alpha}\Big){}_1F_1\Big(1-\frac{Dk^2}{2\alpha};2;\frac{\alpha R^2}{2D}\Big)\\
&-c_1\,\alpha R\,e^{-\frac{\alpha R^2}{2D}}\,{}_1F_1\Big(-\frac{Dk^2}{2\alpha};1;\frac{\alpha R^2}{2D}\Big)\\
&+c_1\,\alpha R\,e^{-\frac{\alpha R^2}{2D}}\,{}_1F_1\Big(-\frac{Dk^2}{2\alpha;1;\frac{\alpha R^2}{2D}}\Big).
\end{aligned}
\tag{S30}
$$

We are then left with the simplified equation,

$$
\Big(\frac{Dk^2}{2\alpha}\Big){}_1F_1\Big(1-\frac{Dk^2}{2\alpha};2;\frac{\alpha R^2}{2D}\Big)=0. \tag{S31}
$$

Here, $k=0$ is satisfied, which yields the steady-state solution. Fig. S10 shows the zeros when we set $\frac{R}{\lambda}=3.16$. The first few non-zero eigenvalues are then $\tilde{k}=0.474$, 0.759, 1.058, 1.354, and 1.672. Here, we observe a similar oscillator pattern to the zeros of the system. Once again, we

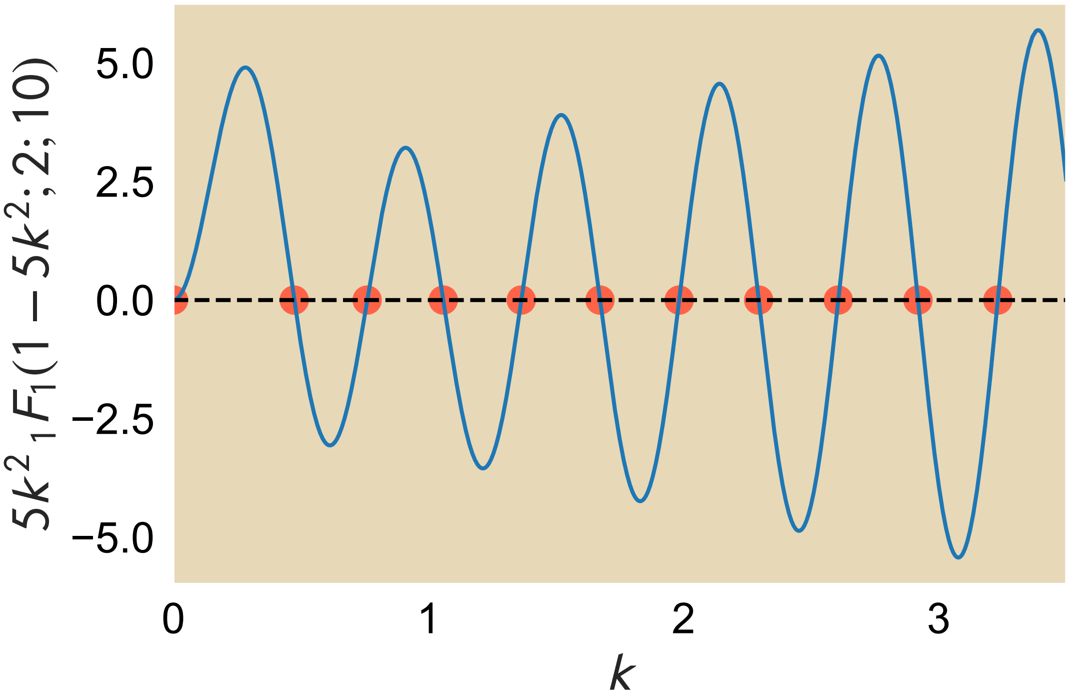

Fig. S10: **Zeros of $k$ for $\frac{\lambda^2k^2}{2}\,{}_1F_1\Big(1-\frac{\lambda^2k^2}{2};2;\frac{R^2}{2\lambda^2}\Big)=0$ where $\frac{R}{\lambda}=3.16$.** Red dots are overlayed with the points where the Kummer confluent hypergeometric function crosses the $x$-axis.

see that there are multiple values of $k$ that satisfy the boundary conditions. This means that the solution to the advection-diffusion problem once both boundary and initial conditions are satisfied, is a superposition of the different eigenfunctions:

$$c(r,t) = c_{\text{ss}}\, e^{-\frac{\alpha r^2}{2D}} + e^{-\frac{\alpha r^2}{2D}} \sum_{i=1}^{\infty} c_i e^{-Dk_i^2 t}\, {}_1F_1\Big(-\frac{Dk_i^2}{2\alpha}; 1; \frac{\alpha r^2}{2D}\Big). \tag{S32}$$

For simplicity, we will reintroduce the length scale $\lambda \equiv \sqrt{\frac{D}{\alpha}}$ so that the equation is simplified as

$$c(r,t) = c_{\text{ss}}\, e^{-\frac{r^2}{2\lambda^2}} + e^{-\frac{r^2}{2\lambda^2}} \sum_{i=1}^{\infty} c_i e^{-Dk_i^2 t}\, {}_1F_1\Big(-\frac{\lambda^2 k_i^2}{2}; 1; \frac{r^2}{2\lambda^2}\Big). \tag{S33}$$

We turn next to the Sturm-Liouville Theory in SI Sec S9 before applying these equations to three simple initial conditions as validation of the theory and the finite element methods (FEM) approach.

### S9 Sturm-Liouville Theory

The Sturm-Liouville theory says that all well-behaved second-order linear ordinary differential equations that can be written in the form

$$\frac{\mathrm{d}}{\mathrm{d}x}\Big[p(x)\frac{\mathrm{d}y}{\mathrm{d}x}\Big] + q(x)\, y(x) = -\lambda\, w(x)\, y(x), \tag{S34}$$

have real eigenvalues with an orthonormal basis of eigenfunctions. Curiously, these equations also have a prescription for determining these eigenfunctions. Importantly, $w(x)$ is the weighting function, which provides the means for satisfying the orthogonality relations for finding coefficients of each term in the series solution to the partial differential equation. Specifically, if we were to write the ODE in the form

$$P(x)\, y''(x) + Q(x)\, y'(x) + R(x)\, y(x) = f(x), \tag{S35}$$

for functions $P(x)$, $Q(x)$, $R(x)$, and $f(x)$, then there is a multiplicative function that can be determined by

$$m(x) = \exp\Big(\int \frac{Q(x) - P'(x)}{P(x)} \mathrm{d}x\Big). \tag{S36}$$

This multiplicative function is then multiplied to Eq. S35 and recast into the form shown in Eq.S34. Thus, with $P(\tilde{x}) = 1$ and $Q(\tilde{x}) = \tilde{x}$,

$$\begin{aligned} m(\tilde{x}) &= \exp\Big(\int \tilde{x}\, \mathrm{d}\tilde{x}\Big), \\ &= \exp\Big(\frac{\tilde{x}^2}{2}\Big), \end{aligned} \tag{S37}$$

and the ODE takes the form

$$0 = \frac{\mathrm{d}}{\mathrm{d}\tilde{x}}\Big[e^{\frac{\tilde{x}^2}{2}} \frac{\mathrm{d}\Phi}{\mathrm{d}\tilde{x}}\Big] + \Phi\Big(1 + \tilde{k}^2\Big) e^{\frac{\tilde{x}^2}{2}}, \tag{S38}$$

or in the form of Eq. S34,

$$\frac{\mathrm{d}}{\mathrm{d}\tilde{x}}\left[e^{\frac{\tilde{x}^2}{2}}\frac{\mathrm{d}\Phi}{\mathrm{d}\tilde{x}}\right]+e^{\frac{\tilde{x}^2}{2}}\Phi=-\tilde{k}^2e^{\frac{\tilde{x}^2}{2}}\Phi, \tag{S39}$$

so that $p(x)=q(x)=w(x)=e^{\frac{\tilde{x}^2}{2}}$ and $\lambda=\tilde{k}^2$.

Next, we show the orthogonality conditions of the eigenfunctions. Suppose that solving Eq. S34 creates a series of eigenfunctions $\{y_j(x)\}$. Suppose that a given eigenfunction $y_i(x)$ has the eigenvalue $\lambda_i$ so that

$$\frac{\mathrm{d}}{\mathrm{d}x}\left[p(x)\frac{\mathrm{d}y_i}{\mathrm{d}x}\right]+q(x)\,y_i(x)=-\lambda_i\,w(x)\,y_i(x). \tag{S40}$$

Suppose that each eigenfunction of the system, bounded by $a\leq x\leq b$, obeys the boundary conditions

$$\begin{aligned}\alpha_1y_i(a)+\alpha_2y_i'(a)&=0,\\ \beta_1y_i(b)+\beta_2y_i'(b)&=0.\end{aligned} \tag{S41}$$

To test the orthogonality conditions, we multiply both sides by $y_j(x)$, a particular eigenfunction of the differential equation, and integrate over the entire system,

$$\begin{aligned}\int_a^b\frac{\mathrm{d}}{\mathrm{d}x}\left[p(x)\frac{\mathrm{d}y_i}{\mathrm{d}x}\right]y_j(x)+q(x)\,y_i(x)\,y_j(x)\mathrm{d}x&=-\lambda_i\int_a^b w(x)\,y_i(x)\,y_j(x)\mathrm{d}x,\\ p(x)\frac{\mathrm{d}y_i}{\mathrm{d}x}y_j(x)\Big|_a^b-\int_a^b p(x)\frac{\mathrm{d}y_i}{\mathrm{d}x}\frac{\mathrm{d}y_j}{\mathrm{d}x}\mathrm{d}x+\int_a^b q(x)\,y_i(x)\,y_j(x)\mathrm{d}x&=-\lambda_i\int_a^b w(x)\,y_i(x)\,y_j(x)\mathrm{d}x.\end{aligned} \tag{S42}$$

Had Eq. S40 involved $y_j(x)$ and we multiplied both sides of the equation by $y_i(x)$, then Eq. S42 would have the subscripts reversed:

$$p(x)\frac{\mathrm{d}y_j}{\mathrm{d}x}y_i(x)\Big|_a^b-\int_a^b p(x)\frac{\mathrm{d}y_i}{\mathrm{d}x}\frac{\mathrm{d}y_j}{\mathrm{d}x}\mathrm{d}x+\int_a^b q(x)\,y_i(x)\,y_j(x)\mathrm{d}x=-\lambda_j\int_a^b w(x)\,y_i(x)\,y_j(x)\mathrm{d}x. \tag{S43}$$

Suppose we subtracted Eq. S43 from Eq. S42 and applied our boundary conditions:

$$\begin{aligned}-(\lambda_i-\lambda_j)\int_a^b w(x)\,y_i(x)\,y_j(x)\mathrm{d}x&=p(x)\frac{\mathrm{d}y_i}{\mathrm{d}x}y_j(x)\Big|_a^b-p(x)\frac{\mathrm{d}y_i}{\mathrm{d}x}y_j(x)\Big|_a^b,\\ -(\lambda_i-\lambda_j)\int_a^b w(x)\,y_i(x)\,y_j(x)\mathrm{d}x&=p(b)\left[\frac{\mathrm{d}y_i}{\mathrm{d}x}\Big|_b y_j(b)-\frac{\mathrm{d}y_j}{\mathrm{d}x}\Big|_b y_i(b)\right]-p(a)\left[\frac{\mathrm{d}y_i}{\mathrm{d}x}\Big|_a y_j(a)-\frac{\mathrm{d}y_j}{\mathrm{d}x}\Big|_a y_i(a)\right],\\ -(\lambda_i-\lambda_j)\int_a^b w(x)\,y_i(x)\,y_j(x)\mathrm{d}x&=p(b)\left[\frac{\beta_1}{\beta_2}y_i(b)\,y_j(b)-\frac{\beta_1}{\beta_2}y_i(b)\,y_j(b)\right]-p(a)\left[\frac{\alpha_1}{\alpha_2}y_i(a)\,y_j(a)-\frac{\alpha_1}{\alpha_2}y_i(a)\,y_j(a)\right],\\ -(\lambda_i-\lambda_j)\int_a^b w(x)\,y_i(x)\,y_j(x)\mathrm{d}x&=0.\end{aligned} \tag{S44}$$

If $i=j$, then the left-hand side is already zero.

$$-\lambda_i\int_a^b w(x)\left[y_i(x)\right]^2\mathrm{d}x=p(x)\frac{\mathrm{d}y_i}{\mathrm{d}x}y_i(x)\Big|_a^b-\int_a^b p(x)\left[\frac{\mathrm{d}y_i}{\mathrm{d}x}\right]^2\mathrm{d}x+\int_a^b q(x)\left[y_i(x)\right]^2\mathrm{d}x. \tag{S45}$$

We will return to the case where $i = j$ to find the coefficients of eigenfunction. If $i \neq j$, then the eigenvalues are different here and the integral is zero:

$$\int_a^b w(x)\, y_i(x)\, y_j(x) \mathrm{d}x = 0, \text{for } i \neq j. \tag{S46}$$

Eq. S45 serves as a convenient equation for analytically solving the coefficients for each eigenfunction.

#### S10 The recovery of a typical FRAP-like disc is time-sensitive in the advection-diffusion model.

As we derive in the SI Sec. S8, the general solution to the PDE

$$\frac{\partial c}{\partial t} = D\nabla^2 c + \nabla \cdot \Big[\alpha \mathbf{r} c\Big], \tag{S47}$$

assuming no angular dependence, takes the form

$$c(r,t) = c_{\mathrm{ss}}\, e^{-\frac{r^2}{2\lambda^2}} + e^{-\frac{r^2}{2\lambda^2}} \sum_{i=1}^{\infty} c_i e^{-Dk_i^2 t}\, {}_1F_1\Big(-\frac{\lambda^2 k_i^2}{2}; 1; \frac{r^2}{2\lambda^2}\Big), \tag{S48}$$

where $c_{\mathrm{ss}}$ is the coefficient for the steady-state concentration term, $\lambda \equiv \sqrt{\frac{D}{\alpha}}$, $k_i$ are the eigenvalues specific to the boundary condition, $c_i$ are the coefficients based on initial conditions, and ${}_1F_1(a;b;z)$ is the Kummer confluent hypergeometric function

$${}_1F_1(a;b;z) = \sum_{l=0}^{\infty} \frac{(a)_l}{(b)_l} \frac{z^l}{l!}, \tag{S49}$$

where the Pochhammer symbol $(a)_l = \frac{(a+l-1)!}{(a-1)!}$. The most well-known example of Eq. S49 is the case where $a = b$, which yields ${}_1F_1(a;a;z) = e^z$. The eigenvalues $\{k_i\}$ are found by satisfying the boundary conditions and are those terms that satisfy the equation

$$\Big(\frac{\lambda^2 k_i^2}{2}\Big) {}_1F_1\Big(1 - \frac{\lambda^2 k_i^2}{2}; 2; \frac{R^2}{2\lambda^2}\Big) = 0. \tag{S50}$$

Eq. S48 shows that the steady-state profile of the concentration is a Gaussian distribution with standard deviation $\lambda$.

We now seek to identify the coefficients of the terms, which are specific to the initial conditions. Here, we will analytically examine three cases for initial conditions: 1) uniform concentration, 2) a uniform concentration except with molecules removed in the region $r \leq R_0$ as found in many FRAP assays, and 3) a FRAP-like removal of molecules in the region $r \leq R_0$ after the system initially reaches a steady-state Gaussian concentration profile. As our goal is to validate our FEM simulations through agreement with some initial conditions that can be analytically determined, we directly compare analytical and FEM solutions.

#### S10.1 Uniform concentration.

We start with the case where the concentration is uniform everywhere,

$$c(r,0) = c_0. \tag{S51}$$

The solution to the PDE with this initial condition takes the form of

$$c(r,t) = \frac{c_0}{2} e^{-\frac{r^2}{2\lambda^2}} \left\{ \frac{\frac{R^2}{\lambda^2}}{1 - e^{-\frac{R^2}{2\lambda^2}}} + \sum_{i=1}^{\infty} \frac{R^2\, e^{-D\,k_i^2 t}\, {}_1F_1\Big(-\frac{\lambda^2 k_i^2}{2}; 2; \frac{R^2}{2\lambda^2}\Big)}{\int_0^R r'\, e^{-\frac{r'^2}{2\lambda^2}} \Big[{}_1F_1\Big(-\frac{\lambda^2 k_i^2}{2}; 1; \frac{r'^2}{2\lambda^2}\Big)\Big]^2 \mathrm{d}r'} \times {}_1F_1\Big(-\frac{\lambda^2 k_i^2}{2}; 1; \frac{r^2}{2\lambda^2}\Big) \right\}. \tag{S52}$$

Fig. S11(A) shows the concentration profile as a function of radius and for various time points given this initial condition. Here, we used $D = 0.1\,\frac{\mu\mathrm{m}^2}{\mathrm{s}}$, $R = 10$ μm, and $v_\mathrm{m} = 0.1\,\frac{\mu\mathrm{m}}{\mathrm{s}}$. Solid lines indicate different time points for the specific analytical solution given the uniform initial condition. These analytical solutions also show strong agreement with simulations performed by FEM which are denoted by hollow points. Here, we use the first 12 eigenvalues $k_i$ for the analytical solution. Similar to the decomposition of a square wave into a sum of sinusoidal functions yielding imperfect agreement with the original function, we see here that the use of a limited number of eigenvalues that satisfy Eq. S50 leads to fluctuations about the original function for $t = 0$ (see SI Sec. S11.6 on Gibbs phenomenon). Nevertheless, we see that these fluctuations in the analytical condition quickly smooth out for $t > 0$. For the given parameters, the concentration at larger radii decreases quickly due to the higher advection overcoming diffusion. As shown at $t = 20$ seconds and $t = 40$ seconds, the concentration appears roughly uniform at lower concentrations but the length scale of this uniformity appears to decrease. At $t = 990$ seconds, the concentration profile reaches the Gaussian steady-state solution where the concentration gradient allows diffusion to counter the advective flow.

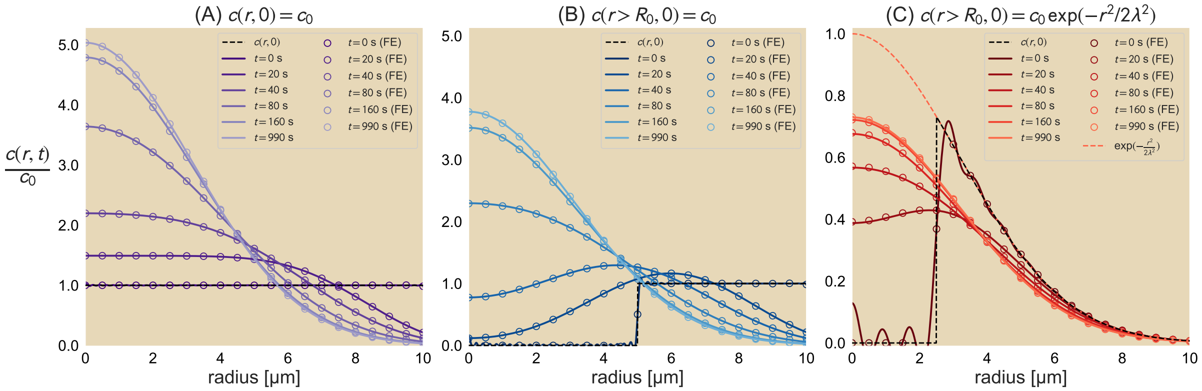

Fig. S11: **Radial advection-diffusion for various initial conditions.** (A) Uniform concentration throughout the system. (B) Uniform concentration for $r > R_0$ and no molecules for $r \le R_0$. (C) A Gaussian distribution for $r > R_0$ and no molecules for $r \le R_0$. Analytical solutions are presented as solid lines while solutions obtained by finite elements are shown as hollow points. The initial condition for each situation is shown as a dashed black line. For all studies, $D = 0.1\,\frac{\mu\mathrm{m}^2}{\mathrm{s}}$, $R = 10$ μm, and $v_\mathrm{m} = 0.1\,\frac{\mu\mathrm{m}}{\mathrm{s}}$. For (B), we set $R_0 = \frac{R}{2}$ while for (C) we set $R_0 = \frac{R}{4}$. For (C), the steady-state profile prior to removing molecules for $r \le R_0$ is shown as a dashed red line. All analytical solutions use the first 12 eigenvalues that satisfy Eq. S50.

##### S10.2 Uniform concentration for $r > R_0$.

We apply a similar initial condition as that used in Sec. S10.1, but remove any molecules within a distance $R_0$ from the origin as typically performed in FRAP experiments. This initial condition is mathematically described by

$$c(r,0) = \begin{cases} 0 & \text{if } r \le R_0, \\ c_0 & \text{if } r > R_0. \end{cases} \tag{S53}$$

The solution for this initial condition is similar to Eq. S52 but with different limits of integration (see SI Sec. S9 on Sturm-Liouville Theory and S8 for application of the theory in 2D),

$$c(r,t) = \frac{c_0}{2} e^{-\frac{r^2}{2\lambda^2}} \left\{ \frac{\frac{R^2}{\lambda^2} - \frac{R_0^2}{\lambda^2}}{1 - e^{-\frac{R^2}{2\lambda^2}}} + \sum_{i=1}^{\infty} \alpha_i e^{-D\,k_i^2 t} \, {}_1F_1\Big( -\frac{\lambda^2 k_i^2}{2}; 1; \frac{r^2}{2\lambda^2} \Big) \right\}, \tag{S54}$$

where

$$\alpha_i = \frac{R^2 \, {}_1F_1\Big( -\frac{\lambda^2 k_i^2}{2}; 2; \frac{R^2}{2\lambda^2} \Big) - R_0^2 \, {}_1F_1\Big( -\frac{\lambda^2 k_i^2}{2}; 2; \frac{R_0^2}{2\lambda^2} \Big)}{\int_0^R r' \, e^{-\frac{r'^2}{2\lambda^2}} \Big[ {}_1F_1\Big( -\frac{\lambda^2 k_i^2}{2}; 1; \frac{r'^2}{2\lambda^2} \Big) \Big]^2 \mathrm{d}r'}. \tag{S55}$$

As $R_0 \to 0$ in Eq. S54 we recover Eq. S52. Fig. S11(B) shows traces of the concentration profile at the same times as in Fig. S11(A). Here, $R_0 = \frac{R}{2}$. Once again, we see that the analytical solution for $t = 0$ fluctuates about the defined initial condition but quickly smooth out and agree well with FEM results (hollow points) for $t > 0$. By removing molecules at $r \le R_0$, a wave of molecules move toward the origin from a combination of advection toward the origin and diffusion moving molecules against the concentration gradient while the concentration at $r \to R$ recedes. Once again, we recover a Gaussian profile, but at a lower maximum than that observed in Fig. S11(A) due to the lower initial number of molecules.

##### S10.3 Gaussian profile for $r > R_0$.

Finally, consider a situation where molecules in this advective-diffusive system are allowed to reach steady-state before photobleaching all molecules within a certain radius of the center $r \le R_0$. The initial conditions would appear as

$$c(r,0) = \begin{cases} 0 & \text{if } r \le R_0, \\ c_0 \, e^{-\frac{r^2}{2\lambda^2}} & \text{if } r > R_0. \end{cases} \tag{S56}$$

We show analytically that the concentration profile is

$$c(r,t) = c_0 e^{-\frac{r^2}{2\lambda^2}} \left\{ \frac{e^{-\frac{R_0^2}{2\lambda^2}} - e^{-\frac{R^2}{2\lambda^2}}}{1 - e^{-\frac{R^2}{2\lambda^2}}} - \frac{1}{2} \sum_{i=1}^{\infty} \beta_i e^{-D k_i^2 t} \, {}_1F_1\Big( -\frac{\lambda^2 k_i^2}{2}; 1; \frac{r^2}{2\lambda^2} \Big) \right\}, \tag{S57}$$

where

$$\beta_i = \frac{R_0^2 \, {}_1F_1\Big( 1 + \frac{\lambda^2 k_i^2}{2}; 2; -\frac{R_0^2}{2\lambda^2} \Big)}{\int_0^R r' \, e^{-\frac{r'^2}{2\lambda^2}} \Big[ {}_1F_1\Big( -\frac{\lambda^2 k_i^2}{2}; 1; \frac{r'^2}{2\lambda^2} \Big) \Big]^2 \mathrm{d}r'}. \tag{S58}$$

Once again the analytical solution agrees with simulations of the same initial condition shown in Fig. S11(C) for $R_0 = \frac{R}{4}$. We note here that as $R_0 \to 0$ we recover the steady-state solution again as the time-dependent terms vanish and the ratio of exponentials in the time-independent term goes to unity. Fig. S11(C) shows again the imperfection of the analytical solution for $t = 0$ and the initial condition but a strong agreement with FEM results. In this situation, the concentration toward the outer edge of the system remains largely unchanged as diffusion and advection are balanced toward the boundary. However, at smaller radii of the system, there is a shift in concentration as molecules enter the $r \leq R_0$ region and for the chosen parameter values, the overall concentration profile returns to a Gaussian distribution within 3 minutes.

Across all three initial conditions, we see that the concentration builds up toward the contraction center and forms a Gaussian distribution as the steady-state profile. The different time courses in the concentration profiles for these initial conditions further reveals that in experimental systems exhibiting such an advective-diffusive behavior the use of FRAP becomes sensitive to the time when photobleaching is applied. If the concentration profile in the system has already begun to move away from a uniform distribution, such as the initial contraction of a highly connected filament network, then the molecule redistribution until steady state is achieved will show different recovery profiles from that of an experiment where photobleaching is applied at a time when the system is already close to reaching the steady-state profile. Such results provide the two extremes of "fluorescence recovery" in potential *in vitro* assays that evolve from a uniform concentration to a Gaussian-shaped distribution subject to this advection-diffusion system.

We show here three cases where analytical solutions to the linear advection-diffusion equation can be determined for direct comparison to the FEM simulations. As the square unit cell is more complex, we turn fully to FEM for our measurements and comparisons to the analyzed experimental data.

### S11 Numerically solving advection-diffusion equations with COMSOL

Our use of COMSOL Multiphysics® simulations are constructed with consideration of four particular details in mind: design of the geometry; set-up of the differential equations, including boundary and initial conditions; choice of mesh size; and sweeping through parameters. Elaboration of the mesh size dependence is discussed in Sec. S11.6.

#### S11.1 Geometry

Because we analyzed fluorescent unit cells from our experimental data until they were no longer distinguishable from neighboring unit cells, we opted to simplify the FEM numerical simulation by examining the time course of a single unit cell subject to advection and diffusion. Even though unit cells in the network may be transported toward the center of contraction, as we have shown in SI Sec S7, the unit cell deformation from advection is not position dependent. This is similarly the case for diffusion, where its contribution to the flux of molecules is dependent on the gradient of concentration. As indicated in Fig. S12, the geometry of the system in the COMSOL simulations is a square of side length 20% longer than the side length of unit cell, which we take to be 10 µm. We then place the smaller square that represents the unit cell inside of the larger square such that it shares the same center. We then take the union of these two squares before applying the split operation to distinguish the unit cell from the surrounding region.

##### S11.2 Setting up the differential equations

Although there are multiple forms of inputting partial differential equations in COMSOL, for the advection-diffusion equation studied, that is

$$\frac{\partial u}{\partial t} = D\nabla^2 u + \alpha\nabla \cdot (\mathbf{r}u)\,, \tag{S59}$$

we elect to use the coefficient form PDE and define our variable of interest as $u$ with units of mol/m$^3$ and ensure that each term in the equation carries units of mol/(m$^3$·s). Although our past derivations use the variable $c$, we use $u$ in the differential equation due to the occurrence of the coefficient $c$ in the coefficient form PDE in COMSOL. We note that the coefficient form PDE as shown in COMSOL is of the form

$$e_a \frac{\partial^2 u}{\partial t^2} + d_a \frac{\partial u}{\partial t} + \nabla \cdot (-c\nabla u - \eta u + \gamma) + \beta \cdot \nabla u + au = f, \tag{S60}$$

where $e_a$, $d_a$, $c$, $a$, and $f$ are scalar coefficients while $\eta$, $\gamma$, and $\beta$ are vectors. We note here that in COMSOL, the term involving $\eta$ is written as $\alpha$, but to avoid confusion with the $\alpha$ used throughout our work, we change the COMSOL notation to $\eta$. Rewriting Eq. S59 to match the form of Eq. S60 gives

$$\frac{\partial u}{\partial t} + \nabla \cdot (-D\nabla u - \alpha\mathbf{r}u) = 0. \tag{S61}$$

We can see here that to make Eq. S61 match Eq. S60, then $e_a$, $a$, all of the elements of $\gamma$, all of the elements of $\beta$, and $f$ are all 0 while

$$d_a = 1\ \mathrm{s}^{-1}, \tag{S62}$$

$$c = D, \tag{S63}$$

$$\eta = \begin{bmatrix} \alpha x \\ \alpha y \end{bmatrix}, \tag{S64}$$

where we note that we define $D$ to take on dimensions of length$^2$/time and $\alpha$ to have units of time$^{-1}$ in COMSOL.

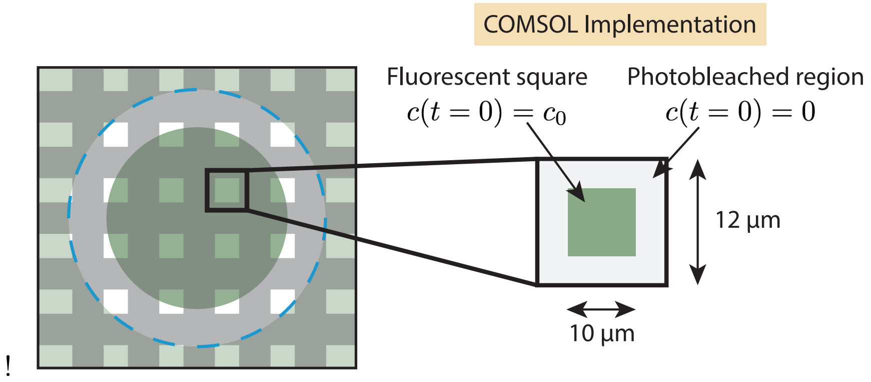

Fig. S12: **Schematic of COMSOL set-up.** To simulate the time evolution of a single unit cell in the advection-diffusion equation, we model a single unit cell as a 10 µm x 10 µm square within a larger 12 µm x 12 µm square.

In our experiments, we are careful to ensure that there is negligible to no detectable amount of microtubules flowing from outside of the light-activated region into network. We thus impose a no-flux boundary condition by using the Zero Flux boundary condition option in COMSOL.

##### S11.3 Applying the initial condition

We opt to make the initial condition of the unit cell of uniform concentration $c_0$ while the concentration in the region outside of the unit cell is initially set to 0. However, defining these two initial conditions piecewise with the geometry of the system outlined above leads to a sharp change in the gradient, which can lead to large errors and negative concentrations at high Péclet number, we instead define a rectangle function where the edges of the rectangle function are smoothed over 200 nm and have well-defined continuous derivatives to second order.

##### S11.4 Choice of mesh size

Because we use the total particle number as a conserved quantity for computing the area of the unit cells in time, we wish to minimize the numerical error in the FEM simulations. Of the various mesh designs, we opt to use the "Extremely fine" mesh size with the boundary between the unit cell and the surrounding system, obtained from the geometry design, to also undergo 6 iterations of refinement under the "Control Entities" tab. This boundary is heavily refined in order to minimize the occurrence and value of negative concentrations that may arise at high Péclet number. A more elaborate discussion of mesh size choice is presented in SI Sec. S11.6 on the Gibbs phenomenon.

##### S11.5 Parameter Sweep

To perform the parameter sweep, we include the Parametric Sweep option in the Study section of the simulation and define the parameters of interest under Global Definitions → Parameters. Within the parameters, we specify the parameters $D$ for our diffusion constant and $\alpha$ for our contraction rate. Under the Parametric Sweep, we can then chose $D$ and $\alpha$ as our parameters to be swept. We select our range of values of $\alpha$ to be the different experimentally-obtained contraction rates while $D$ ranged from 0.0001 $\mu m^2/s$ to 0.01 $\mu m^2/s$ in various increments ranging from 0.0001 $\mu m^2/s$ to 0.0005 $\mu m^2/s$. All possible combinations of $D$ and $\alpha$ were permitted for the simulations.

##### S11.6 Gibbs phenomenon in analytical solutions and mesh granularity in FEM

As we noted in the SI Sec. S10, upon solving the analytical solutions for three cases, there was notable discrepancy between the analytically solved concentration profile at $t = 0$ and the defined initial condition. In this section, we address the sensitivity of the analytical solutions to the number of terms in the infinite series that are kept when showing the concentration profile over time. We then discuss a similar case of sensitivities in the finite element method (FEM) which can also affect the accuracy of numerical solutions.

As shown in Fig. S13, the analytical solution, which is composed of the first 100 non-zero eigenvalues for the two cases involving a uniform initial concentration and the first 25 eigenvalues for the one involving the FRAPed Gaussian profile and the steady-state function, creates oscillations about the intended initial condition. This disagreement is a demonstration of the Gibbs phenomenon, as famously revealed by the imperfect decomposition of a square wave into a sum of sinusoidal functions. Fig. S13 demonstrates the evolution of each of the three analytical solutions examined in the

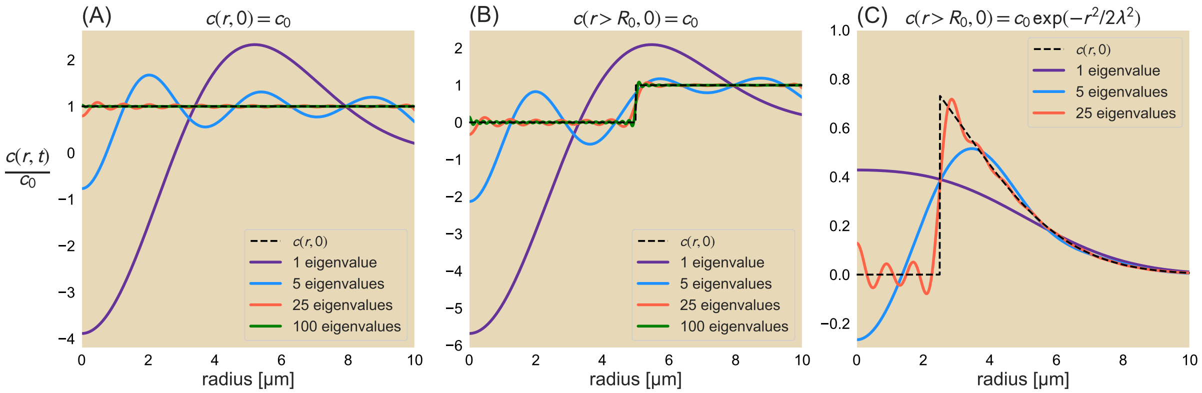

Fig. S13: Gibbs phenomenon for analytical solutions. Concentration profiles of the analytical solution for the initial conditions (A) $c(r, 0) = c_0$, (B) $c(r > R_0, 0) = c_0$, and (C) $c(r > R_0, 0) = c_0\exp(-r^2/2\lambda^2)$ with the steady-state solution and the first nonzero eigenvalue solution (purple line), the first five nonzero eigenvalue solutions (blue), the first twenty-five terms (red), and for (A) and (B) the first hundred terms (green). The intended initial conditions are represented as dashed black lines.

main manuscript when more eigenvalues are included in the solution. Specifically, for $c(r, 0) = c_0$ (Fig. S13(A)), $c(r > R_0, 0) = c_0$ (Fig. S13(B)), and $c(r > R_0, 0) = c_0\exp(-r^2/2\lambda^2)$ (Fig. S13(C)), all of which are represented by dashed black lines, more eigenvalues reduce the level of error between the analytical solution and the initial condition. For the two initial conditions involving a uniform concentration, the use of one eigenvalue in addition to the steady-state solution (purple line) leads to a large negative concentration at $r = 0$ but more closely recapitulate the initial conditions after using 100 non-zero eigenvalues. Deviations from the initial condition decrease dramatically by that point. This is similarly observed for the clipped Gaussian distribution: while the Gaussian tail is quantitatively captured by the the addition of only a few eigenvalues, the analytical solution begins to better recapitulate the concentration profile about $r = R_0$ with the addition of more terms in the solution. Curiously, after using more than 25 eigenvalues, the solution shows large oscillations rather than smaller ones that are smoothed out rather quickly after a small amount of time.

Just as analytical solutions are sensitive to a form of resolution to properly capture the time evolution of a variable of interest, more concretely shown through the number of eigenvalues computed and by extension the number of terms used in the infinite series, so too are there sensitivities in the FEM solution. These sensitivities must also be addressed during setup of the FEM solution to ensure that the model equation is being accurately recapitulated. In this case, a key consideration is the choice of granularity in the mesh. As FEM involves solving the governing equation over a particular domain, having a very fine grained mesh allows for the FEM solution to more accurately reflect the true solution to the problem at the cost of computational time. On the other hand, a very coarse-grained mesh involves less computing power to solve the original equations but may coarse grain away details smaller than the element size, requiring a balance between accurately solving the original PDE(s) and computational efficiency.

Fig. S14 shows how the granularity of the mesh affects the FEM solutions. We compare the concentration profiles produced by FEM (solid blue lines) against the true initial condition (dashed black lines) for six different element sizes as found in the physics-controlled mesh feature in COM-

SOL Multiphysics: (A) extremely coarse, (B) coarse, (C) normal, (D) fine, (E) extra fine, and (F) extremely fine. We see that while using the most coarse-grained feature poorly matches the desired initial condition with more of a sine wave than a square wave, successively decreasing element size (increase in mesh fineness) allows the FEM solution to more closely reflect the initial condition. Fig. S14(B)-(E) show that increasing the mesh fineness leaves fewer deviations from the true values, largely located near the discontinuities in the profile. The insets in the upper right of each figure shows the mesh pattern for the geometry for the study. As Fig. S14(F) shows, while the extremely fine mesh does not overshoot above the $c_0$ values or undershoot the $c(r, 0) = 0$ regions, the finite size of the elements in the mesh causes the discontinuous region to take on a value between the two regions instead. As the FEM simulation is not computationally demanding for the single unit cell case, we opt to use an Extremely Fine mesh setting.

#### S12 FEM results of advection-diffusion equation on a simulated unit cell array

In the main manuscript, we measure the area of the fluorescent squares over time and compare the results to numerical simulations of an advection-diffusion equation through the FEM simulations

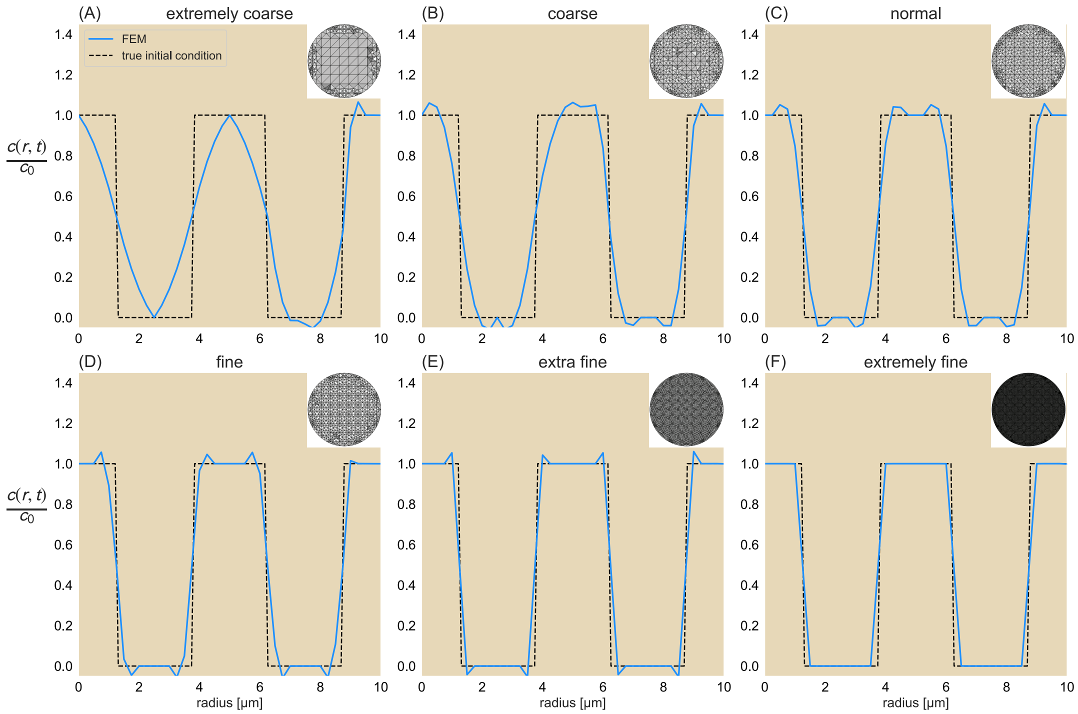

Fig. S14: Effects of mesh granularity on FEM solution. Concentration profiles at $t = 0$ for six different element sizes as defined by the COMSOL Multiphysics physics-controlled mesh: (A) extremely coarse, (B) coarse, (C) normal, (D) fine, (E) extra fine, and (F) extremely fine. Finite elements output is represented by the blues lines while the true initial conditions are given as the black dashed lines. For visualization purposes, the appearance of the meshes used for the defined geometry are shown as insets in the upper righthand corner of the respective subfigures. Concentration profile is from a line trace along the horizontal axis from the origin of the geometry to the boundary.

as described in SI Sec S11 in order to compute effective diffusion constants. While alternative approaches to obtaining the effective diffusion constant exist, we offer this as a direct comparison to numerical experiments. For a qualitative comparison to the experimentally-observed change in the photobleached microtubule network, we present in this section the time evolution of the concentration distribution for an array of unit cells subject to linear advective and diffusive effects. For these simulations, we follow a similar procedure as outlined in SI Sec. S11 but on a circle of radius 60 µm and squares of side length 15 µm with a periodicity of 30 µm. Fig. S15 shows different time points of the concentration profile subject to the same rate of advection (0.002 $\text{sec}^{-1}$) but different diffusion constants, namely, those measured for the median (Fig. S15(A)) and 3rd quartile area trajectories (Fig. S15(B)), an order of magnitude greater diffusion constants (Fig. S15(C)-(D)), and roughly the diffusion coefficient of a free microtubule (Fig. S15(E)). From examining the different concentration profiles in Fig. S15 in comparison to the experimental results shown in Fig. 2 of the manuscript, we see that once again, introducing diffusion to the system is a necessary component to recapitulate the experimentally obtained results. We further see that by eye the simulated data and experiments look most similar when simulating with an effective diffusion constant of 0.1 $\frac{\mu\text{m}^2}{\text{sec}}$. However, we note that as discussed in the results section "The effective diffusion constant is roughly two orders of magnitude lower than free diffusion of a microtubule" in the manuscript and as revealed through Fig. 3, we see that when we careful quantify the area trajectories using the same metric for experiments, the diffusion constants above about $6.0 \times 10^{-3}$ $\frac{\mu\text{m}^2}{\text{sec}}$ would lead to increasing area trajectories instead.

### S13 Motor Constructs

| Motor Construct | Sequence Layout |
|---|---|
| micro variant | pBiex-1:FLAG-GG-mVenus-$(GSG)_2$-micro-$(GSG)_4$-Ncd281 |
| iLid variant | pBiex-1:FLAG-GG-mVenus-$(GSG)_2$-iLid-$(GSG)_4$-Ncd281 |

Table S1: **Ncd281 construct design.** All constructs are designed in the pBiex-1 vector and produced by Twist Biosciences.

| Motor Species | Maximum speed | Processivity |
|---|---|---|
| Ncd281 | 90 nm/s [14] | Nonprocessive [14] |
| Ncd236 | 115 nm/s [13] | Nonprocessive [13, 17] |
| bacterial-expressed K401 | 250 nm/s [1] | ≈ 100 steps [13] |
| insect-expressed K401 | 600 nm/s [13] | ≈ 100 steps [13] (Assumed to be the same as bacterial-expressed K401) |

Table S2: **Motor variant parameters.**

While several of the motors used here in the analysis are obtained from previous work, including K401 expressed in bacteria [1], K401 expressed in insects and Ncd236 expressed in insects [13], we also designed constructs for the study of Ncd281 [14]. Specifically, the sequences are inserted into pBiex-1 vectors and includes a FLAG tag for protein purification, mVenus for motor fluorescence visualization, either a micro or iLid domain as described in [15] and Ncd281 as described in [14]. Between these different domains are multiple repeats of a 'GSG' amino acid sequence which offers flexible links between the regions. Table S1 illustrates these sequences. Constructs were produced

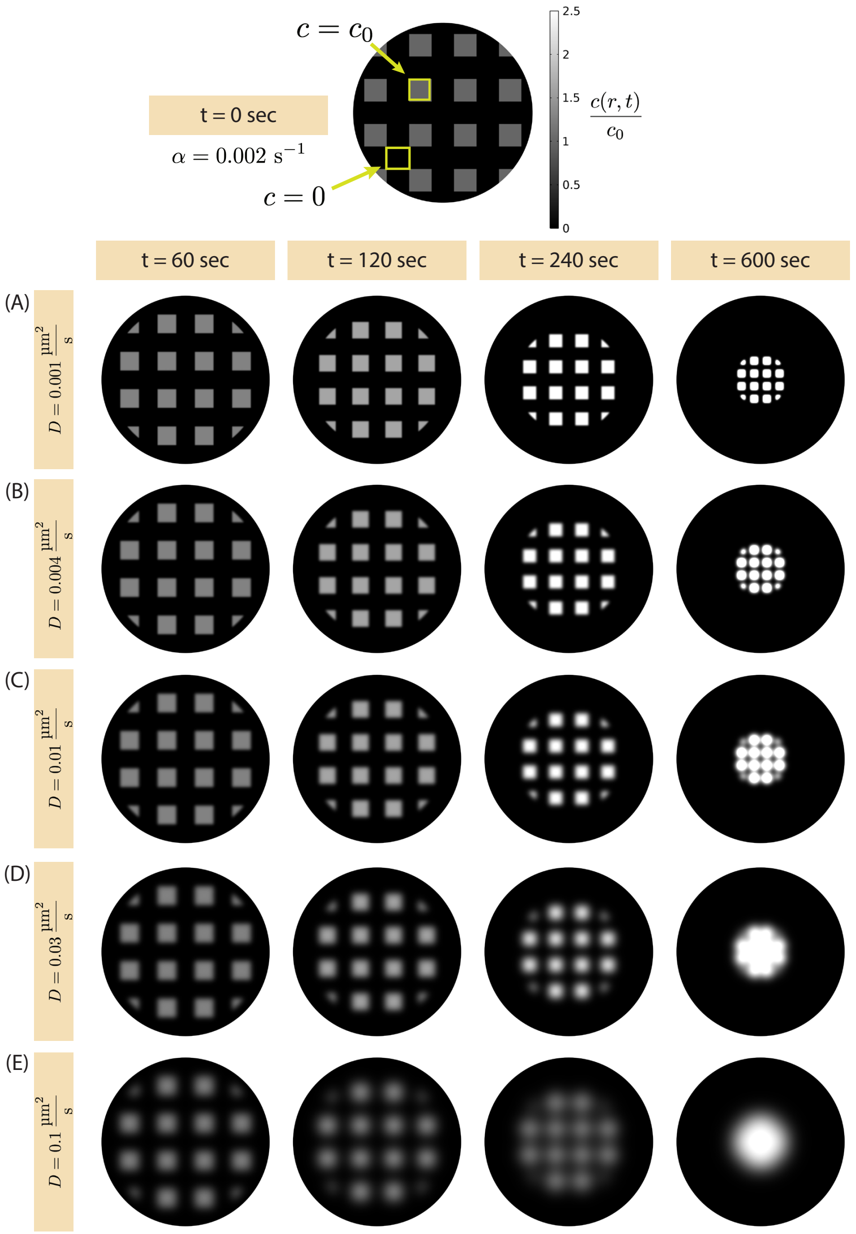

Fig. S15: **Concentration profiles of an array of unit cells at various time points and diffusion constants.** The FEM simulation is the same as that described in SI Sec S11 but where each square (denoted by initial concentration $c_0$ as drawn with the top yellow box in the $t = 0$ sec schematic) has a side length of 15 µm and a center-to-center distance of 30 µm, with a concentration of 0 in between. In all cases, we use the same advection rate of 0.002 sec$^{-1}$ and different diffusion coefficients: (A) 0.001 $\frac{\mu m^2}{sec}$, (B) 0.004 $\frac{\mu m^2}{sec}$, (C) 0.01 $\frac{\mu m^2}{sec}$, (D) 0.03 $\frac{\mu m^2}{sec}$, and (E) 0.1 $\frac{\mu m^2}{sec}$.

by Twist Biosciences.

In addition, Table S2 shows the different motors presented in the manuscript, including their processivities and maximum speeds. Citations of their speeds are added as necessary.

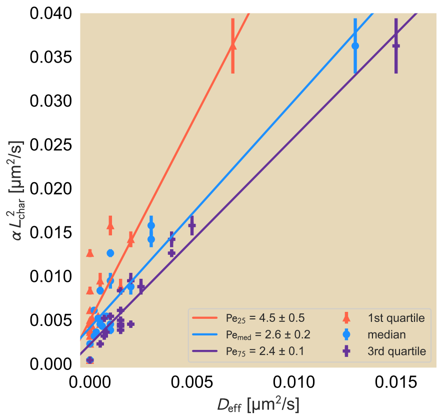

Fig. S16: **Fits of Peclét number for the first quartile (red), median (blue), and third quartile (purple).** We remind the reader that the blue datasets are identical to what is presented in Fig. 6 of the main manuscript.

#### S14 Variability in Peclét number

In the main manuscript, we argue that a Peclét number Pe emerges regardless of the effective motor speed, tuned through ATP concentration or motor species. There, we presented this using the effective diffusion constants fitted from simuations onto the median area trajectories for these conditions. To get an idea for how sensitive Pe is to the variability found within conditions, e.g. the spread in area trajectory distribution, we compute Pe for the first and third quartiles. Fig. S16 shows best linear fits for each of quartiles examined where the slopes denote the respective values of Pe. Here, we find that in addition to the median Peclét number $\mathrm{Pe}_{\mathrm{med}} = 2.6 \pm 0.2$ as noted in the main manuscript, $\mathrm{Pe}_{25} = 4.5 \pm 0.5$ and $\mathrm{Pe}_{75} = 2.4 \pm 0.1$. Here, we see that despite the variability in the effective diffusion constant, Pe is less than a factor of 2 different between the quartiles, suggesting low variability in this dimensionless number.

#### S15 Computing depletion forces

One of the most useful effects of crowding agents is their ability to induce entropic forces upon larger objects when these crowded objects are within the size of the crowding agent from each other. This may be relevant in *in vitro* active systems where the use of crowding agents help to bundle microtubules and promote self-organization. In the case of the work presented here, pluronic ($\backsim$ 12.5 kDa) acts as a crowding agent for microtubules (α and β tubulin have sizes of 50 kDa each and a one-micron long microtubule consists of $\backsim 1.6 \times 10^3$ tubulin). Here, to get a sense of the size of these forces, we estimate the entropic forces induced by crowders such as pluronic onto rigid polymers such as microtubules. To start, we compute the free energy change of the space that crowding agents can occupy when there are two larger particles of radius $R$. We will start by solving in two dimensions where we account only for the cross-sectional area of the microtubules. We will further assume the system does not contain a high density of crowders, so we will say that there are $N$ crowders that can be distributed across $\Omega \gg N$ lattice sites of size $a$. Finally, each crowder will have radius $r$. In the absence of the microtubules, the free energy of the crowders in

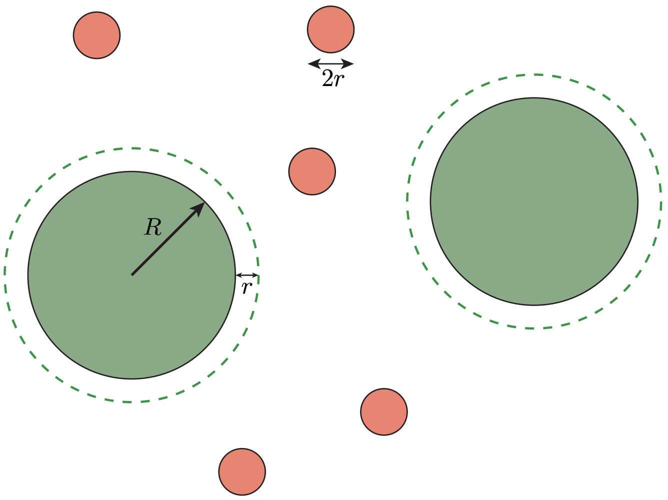

Fig. S17: **Schematic of crowding action on two larger objects.** The crowders (red) have radius $r$ while the two larger objects (green) have radius $R \gg r$. An additional zone around the large molecules as denoted with a dashed outline extends $r$ away from the edge of each molecule and denotes the region within which the centers of the crowders cannot enter.

a system of size $A_{\rm sys}$ is

$$G_{\rm open} = -Nk_BT \ln\left(\frac{A_{\rm sys}}{a}\right), \tag{S65}$$

where $k_BT$ is the thermal energy. Later on we will attempt a derivation where the number of crowders is dense enough where we need to account for their finite size. With the addition of two microtubules, the free energy $G_{\rm crowd}$ becomes

$$G_{\rm crowd} = -Nk_BT \ln\left(\frac{A_{\rm sys} - A_{\rm exc}}{a}\right), \tag{S66}$$

where $A_{\rm exc}$ is the excluded area unavailable to the crowders. This can be represented as the cross-sectional areas of the microtubules with an additional radial buffer zone of length $r$ and depends upon the distance the two cross-sectional areas are from each other. For now, we can compute the free energy change as

$$\begin{aligned}\Delta G \equiv G_{\rm crowd} - G_{\rm open} &= -Nk_BT \ln\left(\frac{A_{\rm sys} - A_{\rm exc}}{A_{\rm sys}}\right), \\ &\approx Nk_BT\,\frac{A_{\rm exc}}{A_{\rm sys}},\end{aligned} \tag{S67}$$

where we assumed that $A_{\rm sys} \gg A_{\rm exc}$.

As noted, the distance between the two microtubules has an effect on the exclusion area. If the microtubules are spaced such that a crowder can fit between them, then $A_{\rm exc}$ is at its maximum, where

$$A_{\rm exc} = 2 \times \pi (R + r)^2. \tag{S68}$$

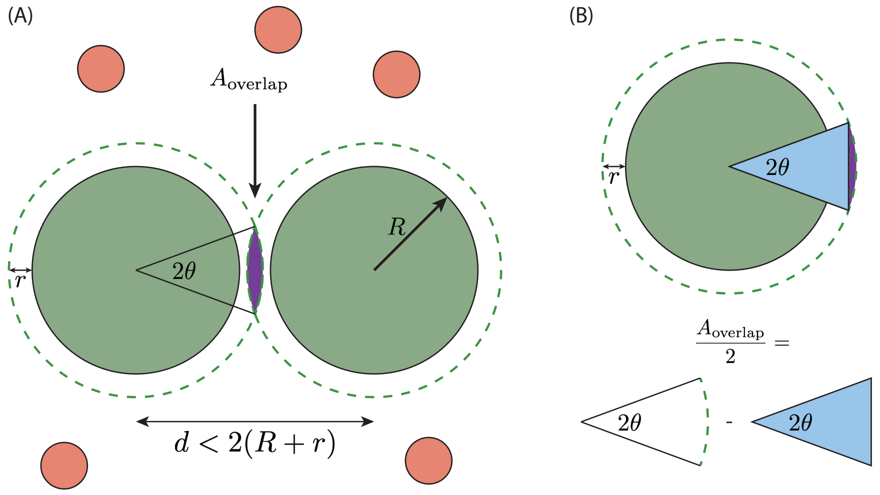

Fig. S18: **Schematic of the overlap of two molecules.** (A) When the large molecules are separated by a distance $d < 2(R+r)$, the exclusion area contains an overlap region that is double-counted in the accounting if the areas of the two molecules and their extended zones are added. (B) The overlap area can be computed by subtracting by computing the difference between the slice of the circle whose arclength begins and ends with the two intersection points of the overlapping circles (as swept out by the angle $2\theta$) and the triangle whose vertices are the center of the circle and the two points where the overlapping circles intersect.

However, if the microtubules are spaced less than a crowder apart, then there is an overlap region that is double-counted $A_{\text{overlap}}$ (Fig. S18(A)). We can compute the area of overlap by recognizing that half of the overlap is the difference between the area swept out by the portion of the circle whose arclength is marked by the intersections of the two overlapping circles and the area of the triangle whose vertices contain these two intersection points and the center of the circle as noted in Fig. S18(B). We will label the common angle between them as $2\theta$.

We can compute the area swept out by the circular slice as

$$\begin{aligned} A_{\text{slice}} &= \int_0^{2\theta} \mathrm{d}\theta' \int_0^{R+r} r' dr', \\ &= \theta(R+r)^2, \end{aligned} \tag{S69}$$

We note that the angle $\theta$ can be obtained with some trigonometry

$$\begin{aligned} \cos\theta &= \frac{d/2}{R+r}, \\ &= \frac{d}{2(R+r)}, \end{aligned} \tag{S70}$$

so the area of the slice as a function of the distance $d$ is

$$A_{\text{slice}} = (R+r)^2 \cos^{-1}\left(\frac{d}{2(R+r)}\right)$$

while the area of the triangle is

$$A_{\text{triangle}} = \frac{d}{2} \times \sqrt{(R+r)^2 - \left(\frac{d}{2}\right)^2}. \tag{S71}$$

Then the overlap region is

$$\begin{aligned} A_{\text{overlap}} &= 2 \times \left(A_{\text{slice}} - A_{\text{triangle}}\right), \\ &= 2(R+r)^2 \cos^{-1}\left(\frac{d}{2(R+r)}\right) - d \times \sqrt{(R+r)^2 - \left(\frac{d}{2}\right)^2}. \end{aligned} \tag{S72}$$

Suppose we made a change of variables to $d = 2(R+r) - \epsilon$ where $0 < \epsilon < 2r$. Then we can modify $A_{\text{overlap}}$ to be

$$\begin{aligned} A_{\text{overlap}} &= 2(R+r)^2 \cos^{-1}\left(\frac{2(R+r)-\epsilon}{2(R+r)}\right) - [2(R+r)-\epsilon] \times \sqrt{(R+r)^2 - \left(\frac{2(R+r)-\epsilon}{2}\right)^2}, \\ &= 2(R+r)^2 \cos^{-1}\left(1 - \frac{\epsilon}{2(R+r)}\right) - [2(R+r)-\epsilon] \times (R+r)\sqrt{1 - \left(\frac{2(R+r)-\epsilon}{2(R+r)}\right)^2}, \\ &\approx 2\,(R+r)^2 \sqrt{\frac{\epsilon}{(R+r)}} - [2(R+r)-\epsilon] \times (R+r)\sqrt{1 - \left(1 - \frac{\epsilon}{2(R+r)}\right)^2}, \\ &\approx 2\,(R+r)^2 \sqrt{\frac{\epsilon}{(R+r)}} - 2\,(R+r)^2 \left(1 - \frac{\epsilon}{2(R+r)}\right)\sqrt{\frac{\epsilon}{(R+r)} - \left[\frac{\epsilon}{2(R+r)}\right]^2} \\ &= 2\,(R+r)^2 \sqrt{\frac{\epsilon}{(R+r)}} \left\{1 - \left(1 - \frac{\epsilon}{2(R+r)}\right)\sqrt{1 - \left[\frac{\epsilon}{4(R+r)}\right]}\right\}, \\ &\approx 2\,(R+r)^2 \sqrt{\frac{\epsilon}{(R+r)}} \left[1 - \left(1 - \frac{\epsilon}{2(R+r)}\right)\left(1 - \frac{\epsilon}{8(R+r)}\right)\right], \\ &\approx 2\,(R+r)^2 \sqrt{\frac{\epsilon}{(R+r)}} \left[\frac{5\epsilon}{8(R+r)}\right], \\ &= \frac{5\,(R+r)^2}{4}\left[\frac{\epsilon}{(R+r)}\right]^{3/2} \end{aligned} \tag{S73}$$

where we note that $\epsilon \ll (R+r)$ and expand to enough orders to maintain a dependence on $\epsilon$. We also note that for small $x$, $\cos^{-1}(1-x) \approx \sqrt{2x}$. As a result, the free energy is

$$\begin{aligned} \Delta G &\approx N k_B T \frac{A_{\text{exc}}}{A_{\text{sys}}}, \\ &= N k_B T \frac{2\pi(R+r)^2 - A_{\text{overlap}}}{A_{\text{sys}}}, \\ &= \frac{N}{A_{\text{sys}}} k_B T \left\{2\pi(R+r)^2 - \frac{5\,(R+r)^2}{4}\left[\frac{\epsilon}{(R+r)}\right]^{3/2}\right\}, \\ &= c k_B T \left\{2\pi(R+r)^2 - \frac{5\,(R+r)^2}{4}\left[\frac{\epsilon}{(R+r)}\right]^{3/2}\right\}, \end{aligned} \tag{S74}$$

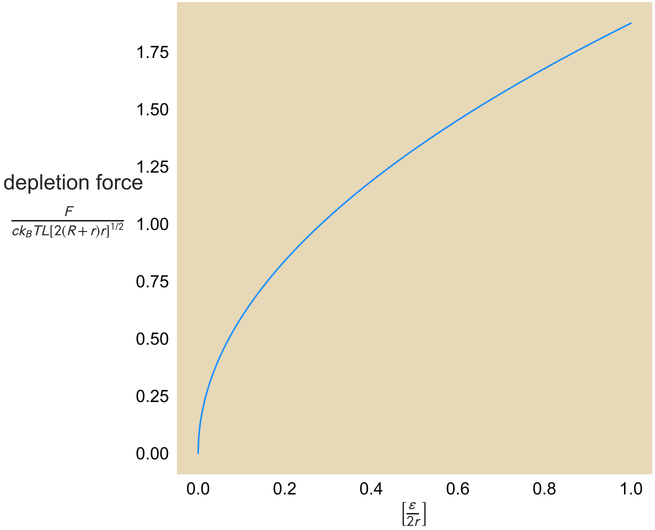

Fig. S19: **Depletion force as a function of overlap distance $\epsilon$.**

where we define the crowder concentration $c = \frac{N}{A_{\text{sys}}}$. As expected, we can see that the free energy goes down as the spacing between the microtubules goes down, suggesting an energetic preference for keeping the microtubules close together.

We can compute the entropic force as the negative derivative of the free energy with respect to the distance $d$. We can then impose the change of variables to see that

$$F_{\text{depletion}} = -\frac{\partial \Delta G}{\partial d} = -\frac{\partial \Delta G}{\partial(2(R+r)-\epsilon)}, \\ = \frac{\partial \Delta G}{\partial \epsilon}, \\ = \frac{15}{8} c k_B T \left[(R+r)\epsilon\right]^{1/2}. \tag{S75}$$

We can then imagine that if we operated in three dimensions, then in the case where two microtubules of length $L$ that are aligned would have a depletion force that goes as

$$F = \frac{15}{8} c k_B T L \left[(R+r)\epsilon\right]^{1/2}. \tag{S76}$$

Fig. S19 shows the relation between the depletion force and the overlap length $\epsilon$. Here, we normalize both sides according to the axes labels. As expected, the depletion force increases as the two microtubules become closer to each other.

If we estimate that a 1 µm-long microtubule has an outer radius of $\backsim$ 10 nm and pluronic, with a mass of 12.5 kDa and a final concentration of 0.5 mg/mL in the experiments (making it 40 µM), has a radius of $\backsim$ 1 nm, then the depletion force due to pluronic is

$$F^{\text{pluronic}}_{\text{depletion}} \backsim \frac{15}{8} \times \frac{40 \times 10^3}{\mu\text{m}^3} \times (4\,\text{pN}\cdot\text{nm}) \times (10\,\text{nm} \times 1\,\text{nm})^{1/2} \times 1\mu\text{m}, \\ \backsim \frac{40 \times 10^4}{\mu\text{m}^2} \times 4\,\text{pN}\cdot\text{nm}^2,$$

$$\backsim 1\,\text{pN}. \tag{S77}$$

Thus, we can see that microtubules under the standard conditions are subjected to roughly picoNewton forces, within the range of forces expected to be exerted by motors. We note, however, that the size of pluronic is even larger, most likely underestimating the computed entropic force.

### S16 Microtubule bundling can affect both contraction speed and filament redistribution

Depletion agents such as pluronic or polyethylene glycol (PEG) play central roles in pushing active systems into contractile or extensile regimes [19, 20]. These polymers help to induce entropic forces between filaments to form bundles, which can help allow active or passive crosslinkers to induce filament coupling over larger length scales. The motor-microtubule system examined thus far includes 0.5 mg/mL pluronic, a concentration that can induce picoNewton-scale forces between microtubules (see SI Sec S7). Here, we ask what happens to the contraction and bulk filament redistribution when these entropic forces are tuned to an alteration to the concentration of pluronic. We thus implement our photobleaching scheme and track the movement and areas of the fluorescent unit cells when the system is altered over a range of pluronic concentrations, from a complete removal of the depletion agent to a 10-fold increase in concentration, while keeping all else fixed, including motor and microtubule concentrations. Fig. S20 shows the contraction rate and effective diffusion constant for the bacterial-expressed K401 across a range of pluronic concentrations, including its complete absence. We find that increasing the pluronic concentration leads to a general increase in the contraction rate until 1.5 mg/mL, after which contraction does not appear to occur any faster. In the absence of pluronic, the network contracts more slowly, with a rate roughly 2/3 the rate of the 1.5 mg/mL pluronic concentration. We note the dramatic decrease at the standard experimental conditions using 0.5 mg/mL pluronic, which lies below even the complete absence of pluronic. We hypothesize that this inconsistency comes from the storage of pluronic in the standard set of experiments being different than the storage conditions used for the pluronic when performing the

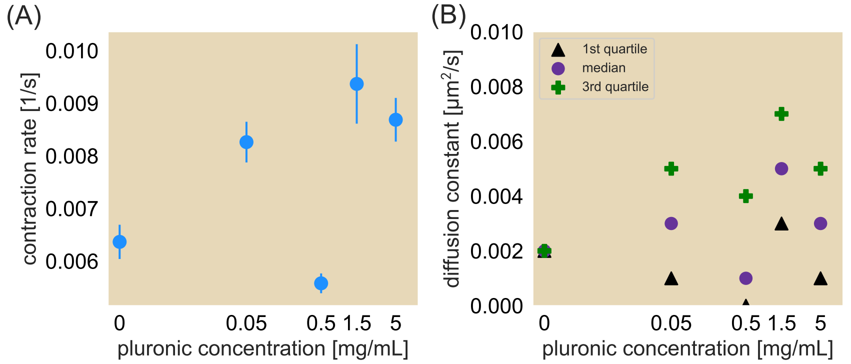

Fig. S20: **The effect of pluronic on the network contraction rate and effective diffusion constant.** (A) Contraction rate and (B) diffusion constant as a function of pluronic concentration as presented here use bacteria-expressed K401 motors. The effective diffusion constants shown here are obtained from best fits to the 1st quartile (triangle), median (circle), and 3rd quartile (plus symbol) of the normalized area trajectories. We note the outlier at 0.5 mg/mL in panel (A) likely corresponds with different storage conditions of the pluronic than from the rest of the other pluronic concentrations used in this study, which may have had a biochemical impact in the assay.

titration series. Briefly, under standard conditions, the pluronic is stored in the base reaction buffer used in the experimental assay involving K-PIPES, $MgCl_2$, EGTA, and KOH. It is possible that under long-term storage in this media, the pluronic behaves differently and as a result exhibits a different effect for the standard reaction.

When we computed the effective diffusion constants for the different quartiles as shown in Fig. S20(B), we found with the increase in pluronic a general increase in effective diffusion constant for the 3rd quartile and median data, but a roughly constant effective diffusion constant for the 1st quartile data. Interestingly, we note that the general increase and decrease of the effective diffusion constants also appears to follow the contraction rate at the corresponding pluronic concentration, suggesting a close relation between the two.

While crowding is commonly implemented in inducing organization in *in vitro* active matter systems and has become a focus of attention as a tunable parameter [21, 22], only recently has crowding been systematically studied to understand its effects on bulk reorganization of a cytoskeletal network [23]. Nevertheless, to our knowledge, we show some of the first experimental studies systematically tuning the effects of crowding on bulk reorganization and observe that entropic forces have more of a binary effect on the contraction rate: in the absence of pluronic, the network contracts more slowly and by adding even 0.1 mg/mL of crowding agent the network contracts more quickly without much more increase in contraction dynamics at higher concentrations. This suggests that entropic forces on the order of pico-Newton scales are sufficient to aid in the formation of a contracted filament network. This is roughly in the same order of magnitude as stall forces for motors, further supporting the role of crowding as generating similar effects to passive crosslinkers.
